# MicroRNA Dynamics and Functions During *Arabidopsis* Embryogenesis

**DOI:** 10.1101/633735

**Authors:** Alexandra Plotnikova, Max J. Kellner, Magdalena Mosiolek, Michael A. Schon, Michael D. Nodine

**Affiliations:** Gregor Mendel Institute (GMI), Austrian Academy of Sciences, Vienna Biocenter (VBC), Dr. Bohr-Gasse 3, 1030 Vienna, Austria; MRC Laboratory of Molecular Biology, University of Cambridge, Cambridge CB2 0QH, United Kingdom

**Keywords:** miRNA, embryo, high-throughput sequencing, RNA-seq, post-transcriptional regulation, transcription factor, *Arabidopsis*, plant development

## Abstract

MicroRNAs (miRNAs) are short non-coding RNAs that mediate the repression of target transcripts in plants and animals. Although miRNAs are required throughout plant development, relatively little is known regarding their embryonic functions. To systematically characterize embryonic miRNAs in *Arabidopsis thaliana,* we developed or applied high-throughput sequencing based methods to profile hundreds of miRNAs and associated targets throughout embryogenesis. We discovered dozens of miRNAs that dynamically cleave and repress target transcripts including 30 that encode transcription factors. Transcriptome analyses indicated that these miRNA:target interactions have a profound impact on embryonic gene expression programs, and we further demonstrated that the miRNA-mediated repression of six transcription factors were individually required for embryo morphogenesis. These data indicate that the miRNA-directed repression of multiple transcription factors is critically important for the establishment of the plant body plan, and provide a foundation to further investigate how miRNAs contribute to these initial cellular differentiation events.

## INTRODUCTION

MicroRNAs (miRNAs) are a class of small regulatory RNAs (sRNAs) that post-transcriptionally repress gene expression and regulate cellular differentiation during plant and animal development (Bartel, 2004; Chen, 2009; Jones-Rhoades et al., 2006; Plasterk, 2006). Plant miRNA precursors fold into characteristic RNA stem-loop structures that are recognized and processed into mature ~21-nt miRNAs by the RNaseIII domain-containing protein DICER-LIKE1 (DCL1) (Park et al., 2002; Reinhart et al., 2002). miRNAs are then loaded onto ARGONAUTE1 (AGO1) proteins and guide the complex to sequences in target RNAs that are almost perfectly complementary to the miRNA (Allen et al., 2005; Jones-Rhoades and Bartel, 2004). In general, miRNAs recognize single sites in target transcripts, and the high-degree of miRNA:target duplex base-pairing results in target RNA cleavage although translational repression has also been reported (Aukerman and Sakai, 2003; Chen, 2004; Gandikota et al., 2007; Jones-Rhoades and Bartel, 2004; Kasschau et al., 2003; Llave et al., 2002). The miRNA-mediated cleavage and repression of transcripts including those encoding transcription factors is required throughout development (Chen, 2009; D’Ario et al., 2017; Jones-Rhoades et al., 2006).

Although miRNAs have been implicated in an array of post-embryonic developmental processes, their functions during embryogenesis remain less well-characterized (Vashisht and Nodine, 2014). This is primarily due to early embryos being small and deeply embedded in maternal seed coat tissues, which makes them difficult to isolate in high purity and characterize the corresponding RNA populations (Schon and Nodine, 2017). Nevertheless, the precursors of the shoot and root meristems, and three main radial tissue layers are precisely established during early embryogenesis and miRNAs are required for most of these early patterning events (Nodine and Bartel, 2010; Schwartz et al., 1994; Seefried et al., 2014; Willmann et al., 2011). Moreover, miRNAs are required to prevent the precocious expression of genes involved in embryo maturation when storage macromolecules such as oil bodies accumulate (Nodine and Bartel, 2010; Willmann et al., 2011). Embryonic miRNAs therefore help define cell-specific gene expression programs according to both spatial and temporal cues. For example, miR165/166 spatially restrict homeobox-leucine zipper family transcription factors during embryogenesis (McConnell et al., 2001; Miyashima et al., 2013; Smith and Long, 2010), and miR156-mediated repression of SQUAMOSA PROMOTER BINDING PROTEIN-LIKE (SPL) transcription factors is required for both the proper divisions of root meristem precursors and to prevent precocious expression of maturation phase genes (Nodine and Bartel, 2010). *mir160a* loss-of-function mutant embryos divide incorrectly, and the abnormal cotyledon phenotypes of seedlings expressing transgenes containing mutations in miR160, miR170/171 or miR319 target sites suggest that the corresponding miRNA activities are required for embryo morphogenesis (Liu et al., 2010; Mallory et al., 2005; Palatnik et al., 2003; Takanashi et al., 2018). The cell-type specific miR394-mediated repression of transcripts encoding the LCR F-box protein is also required for patterning embryonic apical domains (Knauer et al., 2013).

Despite the individual examples of embryonic miRNA functions described above and miRNA profiling studies on late-staged plant embryos (Huang et al., 2013; Oh et al., 2008; Xu et al., 2018), a comprehensive understanding of embryonic miRNA populations and their individual contributions to embryogenesis is incomplete. *Arabidopsis thaliana (Arabidopsis)* embryos are ideal model systems to investigate the roles of miRNAs during plant embryogenesis. Not only do the available genomic and genetic resources in *Arabidopsis* facilitate miRNA functional characterization, but *Arabidopsis* embryos undergo a series of highly stereotypical cell divisions to generate the basic body plan (Mansfield and Briarty, 1991; Palovaara et al., 2016). Therefore, abnormal cell division patterns in early *Arabidopsis* embryos can be screened for upon disrupting miRNA functions in order to test whether the miRNA activities are required for morphogenesis, and thus yield insights into the molecular basis of the corresponding patterning events. In this study, we optimized a low-input small RNA sequencing (sRNA-seq) method to generate profiles of hundreds of miRNAs and used the recently developed nanoPARE approach (Schon et al., 2018) to identify corresponding target transcripts throughout embryogenesis. We found that miRNAs dynamically cleave and repress at least 59 transcripts including 30 encoding transcription factors belonging to eight different families. As a proof-of-principle of this dataset’s utility, we selected individual miRNA/target interactions to investigate further and demonstrated that the miRNA-mediated repression of six transcription factors were individually required for the proper cell division patterns of various post-embryonic tissue-type precursors. Therefore, this resource provides a foundation to further investigate how miRNAs help coordinate the formation of the basic body plan by post-transcriptionally restricting their targets, including transcription factors, to specific stages and cell-types.

## RESULTS

### Establishment of Low-input Small RNA Sequencing Method

To systematically characterize the dynamics and functions of individual embryonic miRNAs in *Arabidopsis,* it was first necessary to determine the miRNAs present in developing embryos. Standard high-throughput sRNA-seq methods however required relatively large amounts of total RNA that were impractical to obtain from early embryos. That is, the sequential ligation of adapters onto the hydroxyl and monophosphate groups at the respective 3’ and 5’ termini of sRNAs followed by reverse transcription and amplification during conventional sRNA-seq library preparation protocols required ≥500 ng of total RNA, which is approximately 100 times more than can be obtained from early *Arabidopsis* embryos. More recent sRNA-seq methods can profile sRNAs from as low as single cells, but do not enrich for sRNAs to the same extent as conventional methods (Faridani et al., 2016). Therefore, to enable the profiling of miRNAs present in developing *Arabidopsis* embryos, we developed a method employing the NEBNext Multiplex Small RNA Library Prep Set for Illumina kit (NEB) to be suitable for the low amounts of total RNA obtainable from early embryos (i.e. 1 – 5 nanograms (ng)). In brief, we included polyacrylamide gel-based size-selection methods to both enrich for sRNAs from total RNA before the first adapter ligation step, as well as to enrich for desired sRNA cDNAs after final PCR amplification (see Methods for details). We also reduced the amounts of 3’ adapters, reverse transcriptase primers and 5’ adapters used in the NEBNext kit when starting with ≤500 ng of total RNA.

We compared sequencing data from libraries generated with 500, 50, 5, 1 or 0.5 ng of total RNA isolated from bent cotyledon stage Col-0 (hereafter referred to as wild-type) embryos to determine how well the method enriches for sRNAs, as well as the method’s reproducibility and accuracy when starting with different amounts of total RNA. Approximately 21-nt miRNAs and 24-nt small interfering RNAs that typically begin respectively with uridine- and adenosine-monophosphates, are characteristic features of plant sRNA populations (Borges and Martienssen, 2015). As expected for plant sRNAs, libraries generated from all input amounts of total RNA predominantly consisted of 21–24-base reads with the first position of the 21- and 24-base reads enriched for thymine and adenine, respectively (Figures 1A and 1B, S1A-S1C). The distribution of sRNA-seq read sizes and 5’ nt biases therefore indicated that the sRNA-seq protocol highly enriches for sRNAs from as low as 0.5 ng of total RNA. To determine the reproducibility of the method across various amounts of input RNA, we compared miRNA family levels between libraries constructed from 500 ng of total RNA with those generated from either 50, 5, 1 or 0.5 ng of total RNA. miRNA levels were highly correlated between biological replicate libraries generated from 500 ng (Pearson’s *R* >0.99) (Figure S1D-S1E). Pearson’s correlation coefficients were >0.9 between 500 ng libraries and all libraries generated with ≥1 ng of total RNA (Figures 1C, S1F-S1H). We also assessed the accuracy of this low-input sRNA-seq method across the dilution series of input RNA by adding exogenous sRNA oligonucleotides (i.e. spike-ins) (Lutzmayer et al., 2017) during RNA isolation prior to library construction, and then examined spike-in levels in the resulting sRNA-seq datasets.

**Figure 1.**
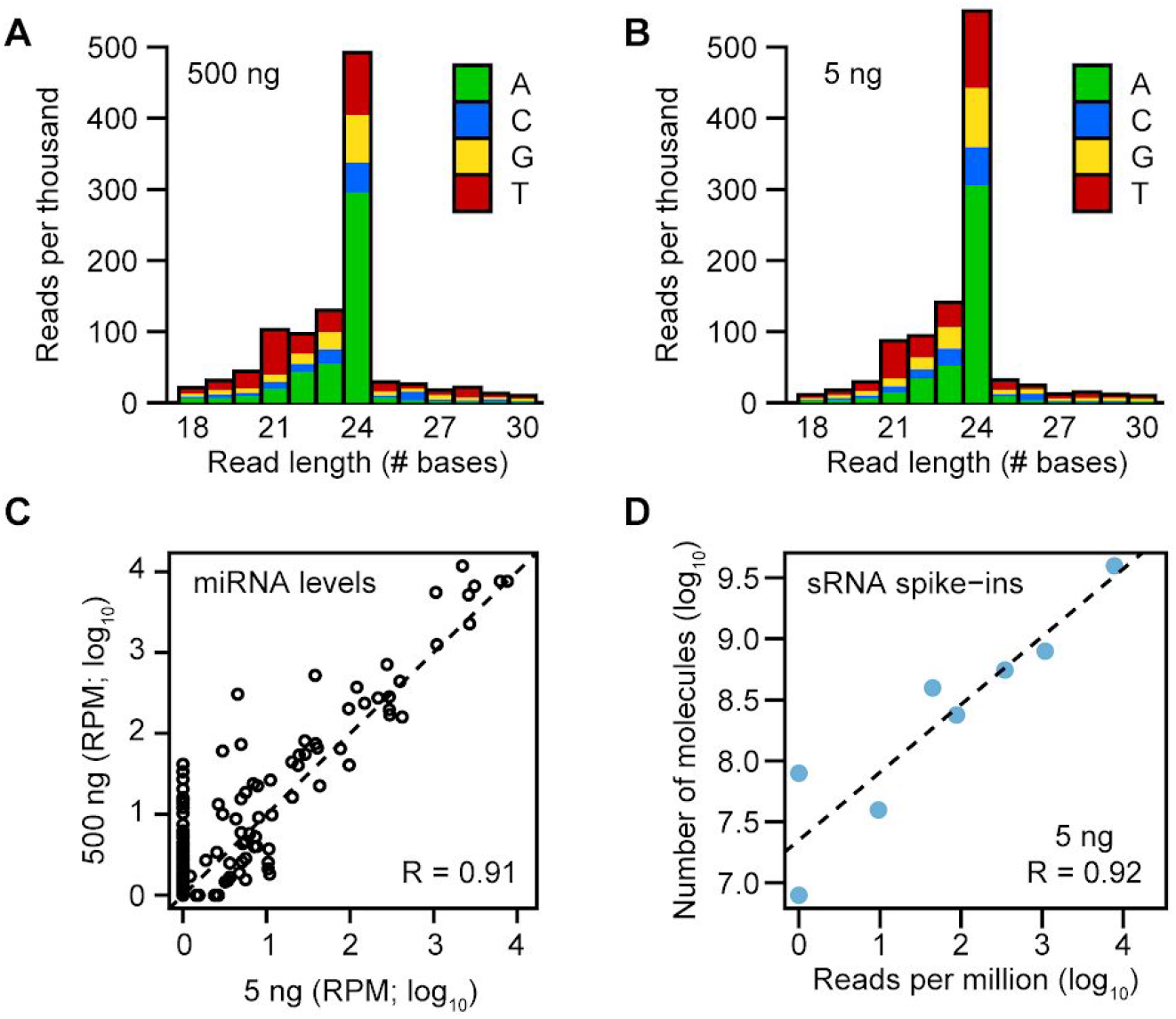
Establishment of Low-Input Small RNA Sequencing Method. (A and B) Stacked bar charts of normalized sRNA-seq read levels (reads per thousand genome-mapping reads) across different base lengths in libraries generated with either 500 ng (A) or 5 ng (B) of total RNA isolated from bent cotyledon stage embryos. Colors indicate the proportions of sRNA-seq reads that begin with various bases as indicated in the key. (C) Scatter plot of miRNA family levels in sRNA-seq libraries generated from 5 ng and 500 ng of total RNA. sRNA levels were normalized for reads per million genome-matching reads (RPM) and log_10_-transformed. Pearson’s *R* value is indicated, as well as a dashed line with an intercept of 0 and slope of 1. (D) Scatter plot of relative sRNA spike-in levels (RPM; log_10_) compared to the absolute number of sRNA spike-in molecules (log_10_) added during RNA isolation for a sRNA-seq library generated from 5 ng of total RNA. Pearson’s *R* value is shown, and the dashed line represents a linear model derived from the plotted data points. See also Figure S1.

If the method accurately quantified sRNA levels, then we expected a high correlation between the absolute number of spike-in molecules added and the number of sRNA-seq reads mapping to the spike-ins. Pearson’s correlation coefficients between the absolute amounts of spike-ins added and the relative amounts of spike-ins sequenced were >0.9 for all libraries generated from ≥1 ng total RNA (Figures 1D and S1I-S1L). The progressive increase in the number of undetected miRNA families and sRNA spike-ins as total RNA amounts decreased indicated that the sensitivity of the method was reduced when starting with less than 50 ng of total RNA (Figures 1C–1D, S1G-S1H, S1K-S1L). Regardless, the modified sRNA-seq library construction method allowed us to highly enrich for sRNAs, and both reproducibly and accurately quantify miRNA levels when starting with 1–5 ng of total RNA, which are amounts obtainable from early *Arabidopsis* embryos.

### Embryonic miRNA Dynamics

We then used this low-input sRNA-seq method to generate libraries with total RNA isolated from embryos at eight developmental stages including three main phases of embryogenesis (Hofmann et al., 2019) (Figure 2A) (Table S1). Three pools of 50 embryos were isolated from each of the eight stages and considered biological replicates (1,200 embryos in total). At least 80% of the total RNA isolated from each biological replicate was used to generate sRNA-seq libraries and the remainder was used to generate full-length cDNAs to profile either transcriptomes (Hofmann et al., 2019) or miRNA-mediated cleavage products (see below). Previous analysis of mRNA-seq libraries generated from the same pools of total RNA demonstrated that the embryonic RNA samples were not significantly contaminated with non-embryonic RNAs, which has been a frequent problem in early embryonic *Arabidopsis* transcriptome datasets (Hofmann et al., 2019; Schon and Nodine, 2017). Total miRNA levels fluctuated in wild-type embryos according to their developmental stage, but were almost completely lost in *dicer-like1-5 (dcl1-5)* null mutants (Figure 2B). Because *DCL1* is required for miRNA biogenesis (Park et al., 2002; Reinhart et al., 2002), this further supports the validity of the miRNAs identified in the sRNA-seq libraries. Principal component analysis of miRNA family levels in libraries generated from embryonic and post-embryonic tissues demonstrated that biological replicates clustered together (Figure 2C). Furthermore, the developmental stages of the embryonic samples were stratified along the second principal component, and were clearly separated from the post-embryonic leaf and flower samples (Figure 2C). By applying the low-input sRNA-seq method to developing embryos, we were therefore able to generate high-quality profiles of embryonic miRNAs, which changed in composition across developmental stages.

**Figure 2.**
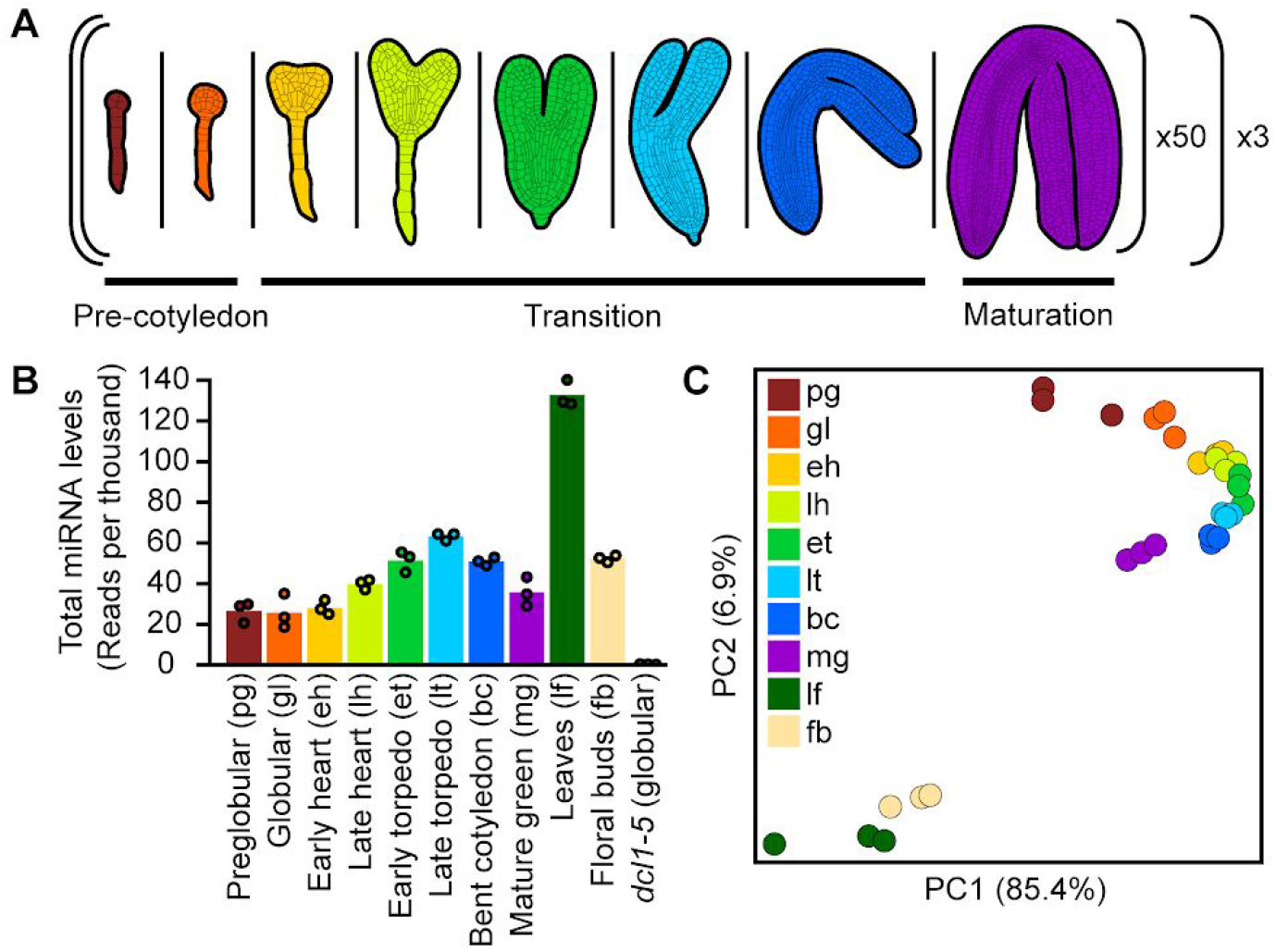
Application of the Low-Input sRNA Sequencing Method to *Arabidopsis* Embryos. (A) Schematic of sRNA profiling experiment across embryogenesis with the low-input sRNA-seq method. Fifty embryos from each of eight different embryonic stages spanning three indicated main phases of embryogenesis were pooled into individual biological replicates. This was repeated three times for each stage to generate three biological replicates for each of the eight developmental stages for (i.e. 24 libraries from a total of 1,200 embryos). (B) Bar chart displaying the total amount of miRNAs detected across wild-type embryogenesis, leaves and floral buds, as well as in miRNA-deficient *dcl1-5* embryos isolated at the globular stage. miRNA levels were normalized by reads per thousand genome-mapping reads. Points indicate the mean levels of individual biological replicates. Individual stages together with their abbreviations are labelled. (C) Principal component analysis illustrating the relationships of the 30 sRNA-seq libraries generated from wild-type embryonic and post-embryonic tissues based on miRNA levels. Embryonic and post-embryonic tissues are labelled according to the key. See also Tables S1 and S2.

We detected 349 miRNAs belonging to 259 families in at least one embryonic stage (Table S2). We then selected 59 miRNA families detected with an average of ≥10 reads per million genome-matching reads (RPM) in at least one embryonic stage to examine in greater detail. Three groups of miRNAs with similar dynamics across embryogenesis were observed (Figure 3A). Twenty-two miRNA families increased during the late transition phase and persisted in mature green embryos. These included miR394, miR403 and miR170/171, as well as miR167 and miR390 which were both previously detected in late-stage embryos with whole-mount in situ hybridizations (Ghosh Dastidar et al., 2016). Another set of 25 miRNA families, including miR156/157, miR161, miR164 and miR319 accumulated during the transition phase but then were reduced in mature embryos. Twelve miRNA families had relatively high levels during early embryogenesis and decreased thereafter. Based on further analysis of internally generated and publicly available sRNA-seq data from 26 tissue-types (Xu et al., 2018), five miRNA families were highly enriched during the initial stages of embryogenesis including miR156b-3p, miR831, miR845, miR866-3p and miR3440b-3p (Figures 3B and S2).

**Figure 3.**
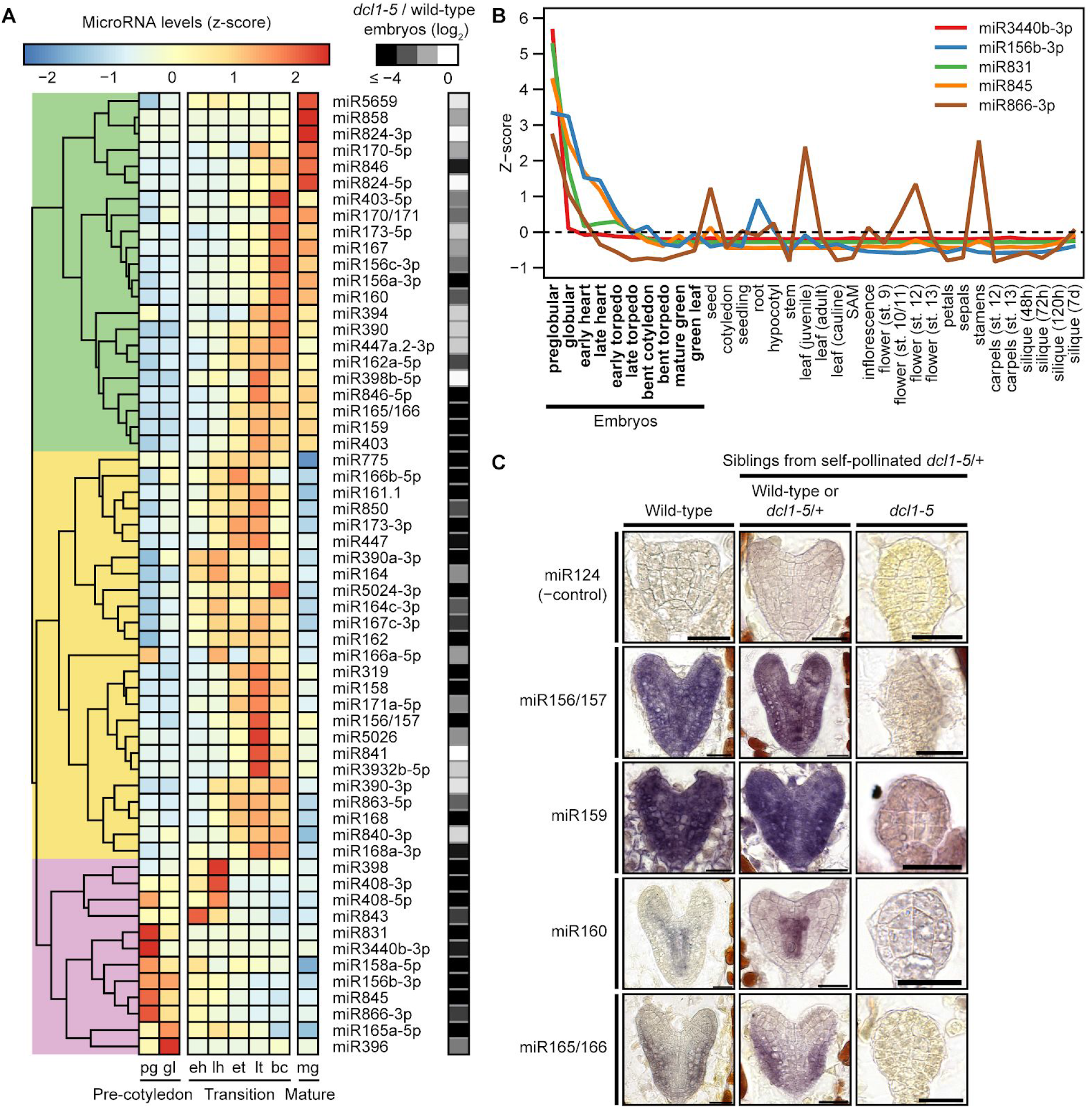
miRNA Dynamics During Embryogenesis. (A) Heat map illustrating the relative levels of miRNA families across embryogenesis. miRNA families with ≥10 mean RPM in at least one embryonic stage are shown, and colors represent z-scores for each individual miRNA family according to the key. Log_2_-transformed levels of miRNAs in (*dcl1-5* + 1)/(wild-type + 1) are annotated. Three major phases of embryo development are labelled at the bottom and individual columns are labelled according to stage: pg, preglobular; gl, globular; eh, early heart; lh, late heart; et, early torpedo; lt, late torpedo; bc, bent cotyledon; mg, mature green. The dendrogram is highlighted in green, yellow or violet to indicate three clusters of miRNA families. (B) Line graph depicting relative levels (z-scores) of preglobular-enriched miRNA families across development. Five miRNA families were selected based on their enrichment in embryos compared to internally generated leaf and floral bud sRNA-seq libraries. sRNA-seq libraries generated internally are marked in bold and published sRNA-seq data from 26 tissue types (Xu et al., 2018) are also shown. (C) Representative images of miRNA in situ hybridizations on sections of embryos. Probes antisense to four miRNAs detected in embryonic sRNA-seq libraries are shown and the corresponding miRNA families are labelled. Probes antisense to the mouse miR124 were used as negative controls, as well as miRNA-deficient *dcl1-5* embryos. Scale bars are 20 μm. See also Figure S2.

To examine whether miRNA levels vary between early embryonic cell-types, we adapted a sRNA in situ protocol (Ghosh Dastidar et al., 2016) to detect four selected miRNAs in early embryos. Consistent with previous reports (Nodine and Bartel, 2010), miR156 was localized throughout wild-type embryos and a similar pattern was also observed for miR159 (Figure 3C). miR165/166 confers repressive activities in the peripheral cell-types of embryos (McConnell et al., 2001; Miyashima et al., 2013; Smith and Long, 2010) and miR166 levels were accordingly higher in these outer cell-types (Figure 3C). In contrast, miR160 levels were higher in the innermost vascular precursor cells (Figure 3C). sRNA in situs performed with probes antisense to the mouse-specific miR124 miRNA produced low signal compared to probes antisense to the four miRNAs in wild-type embryos or embryos with wild-type morphologies from *dcl1-5/+* self-pollinated plants (i.e. wild-type or *dcl1-5/+* embryos) (Figure 3C). Moreover, probes antisense to the four miRNAs produced highly decreased signals when applied to miRNA-deficient *dcl1-5* embryos compared to wild-type or *dcl1-5/+* embryos (Figure 3C). These controls further support the specificity of the signal observed from the miRNA in situ hybridizations.

### Identification of Embryonic miRNA Targets

Based on our analyses, embryonic miRNA populations were distinct from those in post-embryonic tissues and their levels frequently exhibited dynamic changes across developmental stages and sometimes cell-types. These results suggested that miRNAs have distinct functions during different phases of embryogenesis. Because miRNA functions are largely defined by the targets they regulate, we next determined the targets of embryonic miRNAs. In plants, miRNAs typically bind to highly complementary binding sites within target RNAs and mediate their endonucleolytic cleavage (Jones-Rhoades and Bartel, 2004; Kasschau et al., 2003; Llave et al., 2002). miRNA-mediated cleavage of target RNAs produces cleavage products downstream of the slice site, which can be cloned and sequenced with high-throughput methods referred to as PARE (parallel analysis of RNA ends) or degradome sequencing (Addo-Quaye et al., 2008; German et al., 2008; Gregory et al., 2008). Although these ground-breaking technologies have allowed miRNA target identification on a genome-wide scale, they required ≥10,000-fold more input RNA than what was obtainable from early *Arabidopsis* embryos. We previously developed a method called nanoPARE to enable the confident identification of miRNA-mediated target RNA cleavage products from low-input RNA (Schon et al., 2018). To identify embryonic miRNA targets, we therefore generated nanoPARE libraries from the same eight stages of embryogenesis used for miRNA profiling in biological triplicates. In addition to these 24 libraries from wild-type embryos, we also generated nanoPARE libraries from three biological replicates of *dcl1-5* globular embryos as controls (Table S1).

The nanoPARE datasets and target predictions for 164 miRNAs detected ≥1 RPM in ≥1 embryonic stage were used as input for the *EndCut* software (Schon et al., 2018) and 115 significant individual target transcript cleavage sites in developing embryos were identified in ≥1 library (Benjamini-Hochberg adjusted *P*-values < 0.05) (Table S3). These 115 target sites included 59 sites that were identified in ≥2 biological replicates from ≥1 developmental stage (i.e. high-confidence targets) for 22 miRNA families that we characterized further. The first positions of nanoPARE reads mark RNA 5’ ends and we found that nanoPARE reads at the 59 high-confidence target sites detected in wild-type embryos were significantly reduced 40.5-fold in miRNA-deficient *dcl1-5* globular embryos (*P*-value < 0.0001; two-tailed K-S test) (Figure 4A). Moreover, no high-confidence targets were detected in *dcl1-5* embryos and 58/59 of the high-confidence targets detected in developing wild-type embryos had decreased nanoPARE reads in *dcl1-5* embryos (Figure 4B). A lack of signal in *dcl1-5* embryos could be explained by either a loss of miRNA-mediated cleavage or technical differences in sample RNA quality. To differentiate between these two explanations, we measured nanoPARE signal mapping to published transcription start sites (Schon et al., 2018) of all high-confidence targets detected in globular embryos. Full-length transcripts were more abundant in *dcl1-5* embryos for 17/20 of these high-confidence targets, demonstrating that the observed reduction of nanoPARE signal at miRNA-directed cleavage sites in *dcl1-5* embryos was not due to differences in RNA quality(Figure S3). The loss of miRNA-mediated cleavage sites in miRNA-deficient *dcl1-5* embryos further supports the validity of the miRNA targets identified.

**Figure 4.**
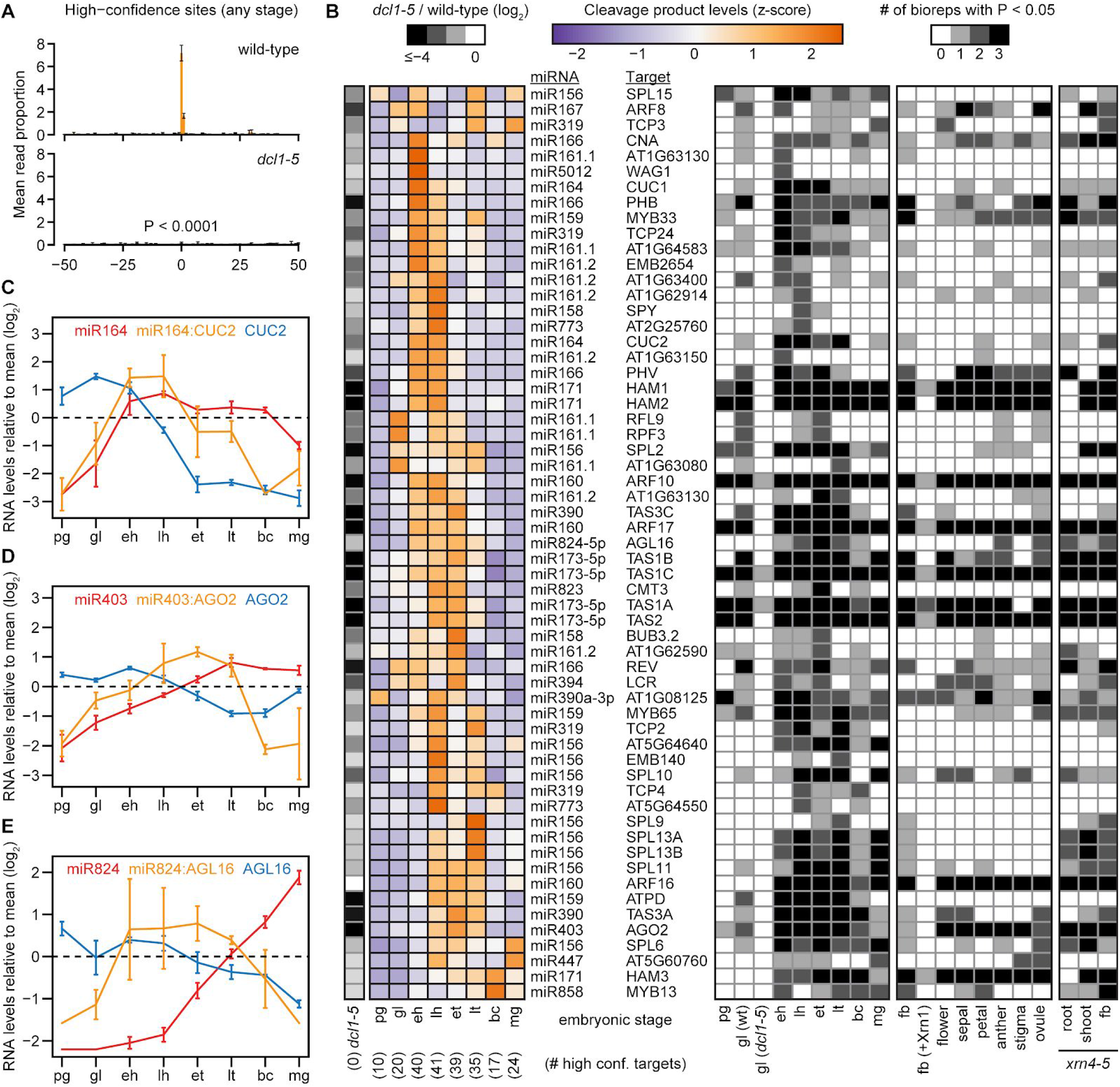
Identification of Embryonic miRNA Targets. (A) The proportion of nanoPARE reads mapping within 50-nt of miRNA target sites significantly detected by *EndCut* (Benjamini-Hochberg adjusted *P*-values < 0.05) in ≥2 biological replicates from any embryonic stage (i.e. high-confidence miRNA cleavage sites; n = 59) are shown for wild-type (top) and *dcl1-5 (bottom’)* globular embryo libraries. The *P*-value that the mean number of reads at the predicted cleavage sites in *dcl1-5* is different than the wild-type mean due to chance is indicated (two-tailed K-S test). Error bars represent the standard errors of the means of three biological replicates. (B) Heat maps depicting the relative levels of miRNA-mediated cleavage products *(left)* and the number of biological replicate libraries in which cleavage products were significantly detected *(right).* High-confidence miRNA cleavages sites from any embryonic stage are shown together with miRNA families and target transcripts alongside the rows. Colors represent z-scores as indicated in the key. Log_2_-transformed levels of cleavage products in *(dcl1-5* + 1)/(wild-type + 1) are annotated, and embryonic stages are indicated below each column, as well as the number of high-confidence targets from that stage. *(right)* Shading densities in the right heatmap indicate the number of biological replicates for which the cleavage site was significantly detected by *EndCut* according to the key (Benjamini-Hochberg adjusted *P* < 0.05). Embryonic stages and post-embryonic tissues in wild-type and *xrn4-5* genotypes are indicated below each column. *dcl1-5*, *dcl1-5* globular; pg, preglobular; gl, globular; eh, early heart; lh, late heart; et, early torpedo; lt, late torpedo; bc, bent cotyledon; mg, mature green; fb, unopened floral buds; +Xrn1, RNA treated with Xrn1 exoribonuclease prior to library construction. (C – E) Line graphs illustrating the relative RNA levels of miRNAs (red), targets (blue) and miRNA-mediated cleavage products (orange) for miR164:CUC2 (C), miR403:AGO2 (D) and miR824:AGL16 (E). Error bars represent the standard errors of the means of three biological replicates for each stage. Relative levels of miRNAs (RPM), cleavage products (reads per ten million genome-matching reads) and transcripts (TPM) for each stage were calculated by log_2_-transforming (stage levels + 1)/(mean levels across all stages + 1). Embryonic stage abbreviations below each graph are as indicated in panel B. See also Table S3.

miRNA-mediated cleavage products dynamically accumulated and were generally more abundantly detected during mid-embryogenesis (Figure 4B). To test whether there are sites enriched in embryos, we also analyzed nanoPARE libraries generated either previously from flowers and floral organs (Schon et al., 2018), or in this study from root or shoot tissues of *xrn4-5* mutants which stabilize miRNA-directed cleavage products (German et al., 2008; Souret et al., 2004). We observed 11 high-confidence target transcripts enriched in developing embryos including those encoding the EMB2654 (miR161.2) and SPY (miR158) tetratricopeptide repeat proteins respectively involved in embryogenesis and gibberellic acid responses, an ATP synthase delta subunit (ATPD; miR159), a plant invertase/methylesterase inhibitor family protein (AT5G64640; miR156/157), and TCP4 and TCP24 (miR319) transcription factors. In fact, 30/59 high-confidence targets encoded transcription factors, and a few cleavage products exhibited interesting developmental dynamics. For example, miR164:CUP-SHAPED COTYLEDON 2 (CUC2) and miR824:AGAMOUS-LIKE 16 (AGL16) cleavage products accumulated during mid-embryogenesis when miRNA and target levels were increasing and decreasing, respectively (Figure 4C and 4E). Similarly, miR403:ARGONAUTE 2 (AGO2) products had relatively high levels during mid-embryogenesis when increasing miR403 levels were concomitant with decreasing AGO2 levels (Figure 4D).

### Impact of miRNAs on the Embryonic Transcriptome

To assess how miRNAs influence embryonic transcript levels, we profiled transcriptomes from *dcl1-5* globular embryos in which miRNA levels and cleavage activities were highly reduced (Figures 2B, 3A, 3C, 4A and 4B). Principal component analysis of *dcl1-5* and wild-type embryonic transcriptomes (Hofmann et al., 2019) revealed that *dcl1-5* biological triplicates clustered together in a group that was separate from the wild-type transcriptomes (Figure 5A). This suggested that the loss of miRNAs resulted in large-scale changes in transcript populations. Indeed, 2,055 and 830 genes had at least two-fold significantly increased and decreased transcript levels in *dcl1-5* relative to wild-type globular embryos, respectively (DESeq2; Benjamini-Hochberg adjusted *P*-values < 0.01) (Figure 5B) (Table S4). Considering that 18,420 genes had ≥1 TPM in either wild-type or *dcl1-5* globular embryos, this indicated that >15% of the transcriptome is significantly altered in *dcl1-5* embryos.

**Figure 5.**
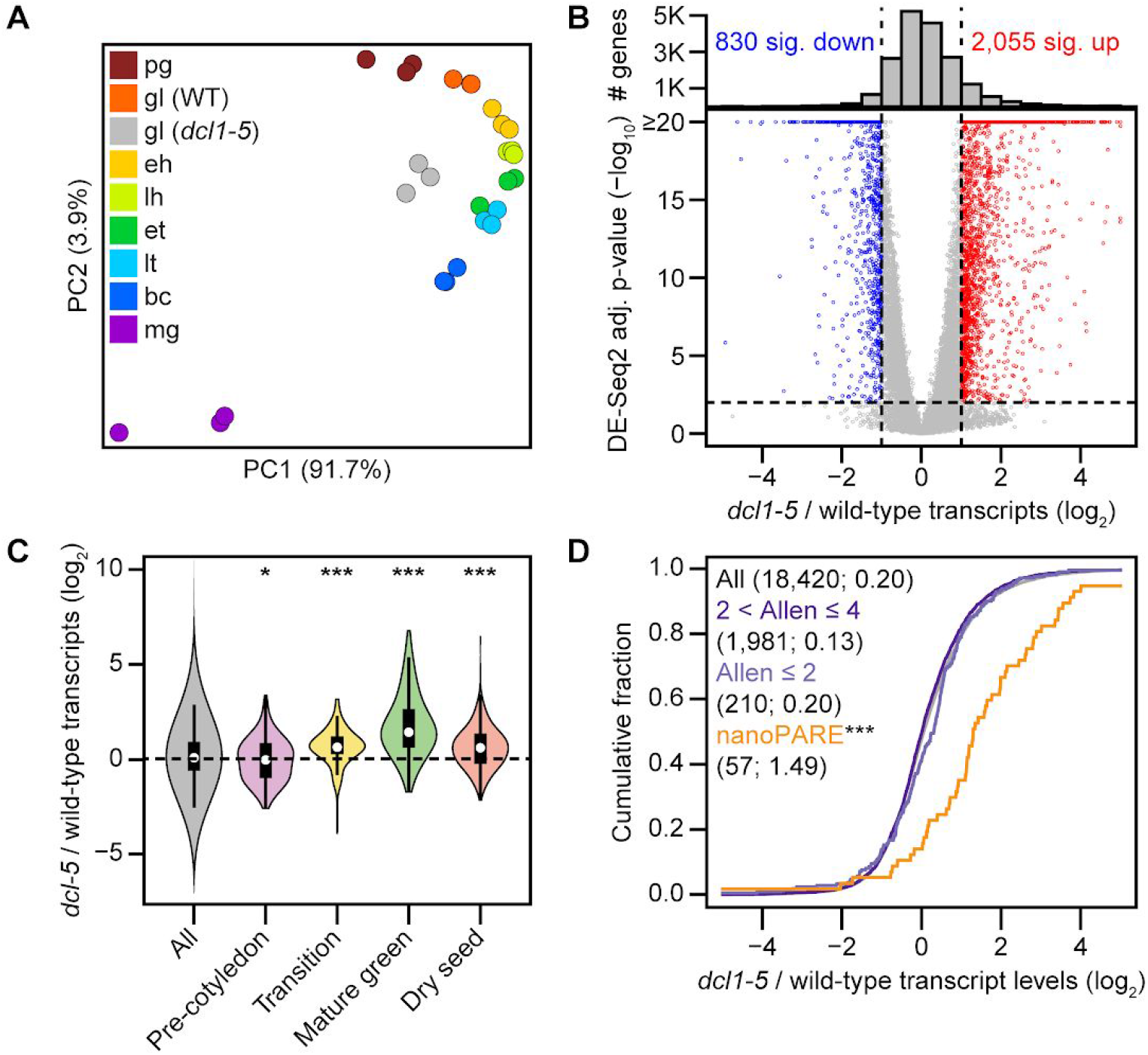
Impact of miRNAs on the Embryonic Transcriptome. (A) Principal component analysis illustrating the relationships between mRNA-seq libraries generated from biological triplicates of either *dcl1-5* globular embryos or wild-type embryos from eight stages. As shown in the key, libraries are color-coded according to their stage: pg, preglobular; gl (WT), wild-type globular; gl *(dcl1-5), dcl1-5* globular; eh, early heart; lh, late heart; et, early torpedo; lt, late torpedo; bc, bent cotyledon; mg, mature green. (B) Volcano plot of log_2_-transformed transcript levels in (*dcl1-5* TPM + 1)/(wild type TPM + 1) (x-axis) and log_10_-transformed Benjamini-Hochberg adjusted *P*-values based on DESeq2 (Love et al., 2014) in *dcl1-5* compared to wild-type (y-axis) *(bottom).* Red and blue indicate transcripts with *P*-values < 0.01 and ≥2-fold increased levels in *dcl1-5* or wild-type embryos, respectively. Histogram of the total numbers of transcripts across different dcl1-5/wild-type transcript fold-changes are shown (top). The number of significantly decreased (sig. dec.) and increased (sig. inc.) transcripts are indicated. (C) Violin plots of log_2_-transformed transcript levels in *(dcl1-5* + 1)/(wild-type + 1) for embryo phase-enriched marker transcripts as defined previously (Hofmann et al., 2019). Transcripts with ≥1 TPM in either *dcl1-5* or wild-type globular embryos (All; n = 18,420) were subset for those enriched in either pre-cotyledon (n = 107), transition (n = 127), mature green (n = 48) or dry seed (n = 183) phases. *P*-values < 0.01 and *P*-values < 10^-6^ are indicated by * and ***, respectively (two-tailed K-S test). (D) Cumulative distributions of *(dcl1-5* TPM + 1)/(wild-type TPM + 1) transcript levels (log_2_) for all transcripts with ≥1 TPM in either *dcl1-5* or wild-type globular embryos (All; black), transcripts confidently predicted computationally (2 < Allen scores ≤ 4, dark purple; Allen scores ≤ 2, light purple), and high-confidence targets detected by *EndCut* (nanoPARE, orange; *** indicates *P*-values = 3.96 × 10^-11^). See also Table S4 and Figure S4.

Differences due to RNA contamination from non-embryonic seed tissues could be ruled out by applying the tissue-enrichment test (Schon and Nodine, 2017), which revealed no significant RNA contamination in either the wild-type or *dcl1-5* samples (Figure S4). These large-scale transcriptome changes may be related to the precocious activation of embryo maturation gene expression programs as previously reported (Nodine and Bartel, 2010; Willmann et al., 2011). To test whether transcriptomes from *dcl1-5* globular embryos resembled those from later stages of development, we examined the levels of transcripts that were specifically enriched during one of four main phases of embryogenesis and seed development (Hofmann et al., 2019) in *dcl1-5* compared to wild-type globular embryos. Transition, mature green and dry seed phase marker transcripts were significantly increased (*P*-values < 10^-6^, two-tailed K-S tests; Figure 5C) indicating that miRNA-deficient *dcl1-5* embryos indeed prematurely activate late-stage gene expression programs.

Because we detected relatively few miRNA targets with nanoPARE compared to the total number of differentially expressed genes (Table S4 and Figures 4B and 5B), we reasoned that either many miRNA targets were not detected with nanoPARE or that the large-scale changes in *dcl1-5* transcriptomes were mostly a consequence of miRNA target de-repression. Plant miRNA targets can be predicted at various confidence levels depending on the frequency and position of the miRNA:target duplex mismatches (i.e. Allen scores) (Allen et al., 2005). Whereas miRNA targets detected with nanoPARE were significantly increased in *dcl1-5* relative to wild-type embryos compared to all expressed genes (2.8-fold; *P*-value = 3.96 × 10^-11^, two-tailed K-S test), targets confidently predicted computationally (i.e. Allen scores ≤ 2 or ≤4), including bona fide post-embryonic targets, were not substantially increased in *dcl1-5* (Figure 5D). These results are consistent with the loss of miRNAs in *dcl1-5* embryos resulting in the loss of cleavage and repression of dozens of targets, and the consequential mis-regulated miRNA target activities having a large impact on embryonic gene expression programs.

Thirty of the fifty-nine high-confidence miRNA targets detected with nanoPARE encoded transcription factors belonging to eight different families including those containing ARF, GRAS, HD-ZIP, MADS-box, MYB, NAC, SBP and TCP domains (Figure 6A). Twenty-eight of these had transcripts >1 TPM in either wild-type or *dcl1-5* globular embryos, and remarkably twenty-four (85.7%) were significantly up-regulated in *dcl1-5* compared to wild-type embryos (Figure 6B). RNA in situ hybridizations of three miR165/166 target RNAs encoding class III HD-ZIP transcription factors (i.e. PHABULOSA (PHB), CORONA (CNA) and PHAVOLUTA (PHV)) in wild-type embryos were congruous with previous reports (McConnell et al., 2001; Prigge et al., 2005; Smith and Long, 2010) and transcriptome analyses (Hofmann et al., 2019). Consistent with their up-regulation in *dcl1-5* embryos, PHB, CNA and PHV transcripts had increased signal throughout embryos including ectopic localization in the basal and peripheral regions of *dcl1-5* embryos (Figure 6C). Together with the observation that miR165/166 and its target transcripts had opposite localization patterns in heart-staged wild-type embryos (Figures 3C and 6C), the ectopic localization of class III HD-ZIP transcripts further supports that miR165/166 helps define the proper localization patterns of their target transcription factors.

**Figure 6.**
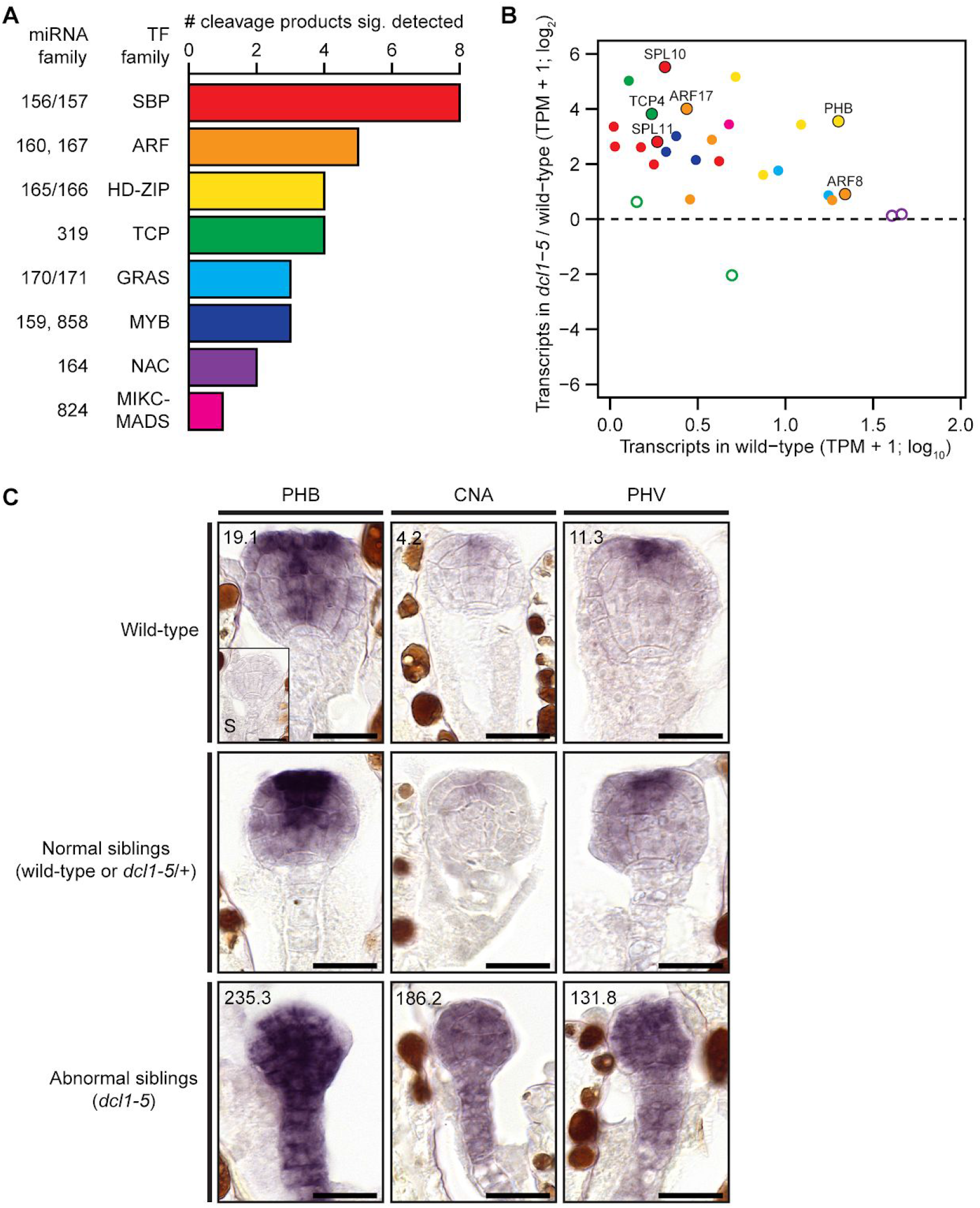
miRNA-mediated Cleavage and Repression of Transcripts Encoding Transcription Factors. (A) Stacked bar chart indicating the number of transcription factor family members for which high-confidence cleavage products were detected during embryogenesis. Transcription factor families, as well as the miRNA families that mediate their cleavage are labelled at the bottom. (B) Scatter plots illustrating the levels of transcripts encoding transcription factors for which high-confidence cleavage products were detected according to their levels in wild-type (TPM + 1; log_10_) and relative fold-changes in log_2_-transformed *(dcl1-5* TPM + 1)/(wild-type TPM + 1). Transcripts with >1 TPM in either wild-type or *dcl1-5* embryos are shown (n = 28). Significantly increased transcripts (n = 24; DESeq2 Benjamini-Hochberg adjusted *P*-values < 0.01; DESeq2) are indicated by points filled with colors representing various transcription factor families as shown in (A). Six targets selected for further analyses are labelled directly above each corresponding point and are also indicated with outlines. (C) Representative images of RNA in situ hybridizations with probes antisense to either PHB *(left),* CNA *(middle’)* or PHV *(right)* transcripts on sections of embryos. RNA in situs were performed on embryos from either self-pollinated wild-type (top) or *dcl1-5/+* plants. Embryos from self-pollinated *dcl1-5/+* plants were further classified into either normal (*middle*, wild-type or *dcl1-5/+)* or abnormal (*bottom*, *dcl1-5)* siblings based on morphology. A sense control for PHB (S) is displayed in the inset of the top left panel. Numbers in top left corners of wild-type and abnormal sibling images are transcript levels (TPMs) determined by mRNA-Seq. Scale bars represent 20 μm.

### miRNA-mediated Repression of Transcription Factors is Required for Embryo Morphogenesis

Embryonic miRNAs direct the cleavage and repression of at least thirty transcripts encoding transcription factors (Figures 4B, 6A and 6B). This appears to help define transcription factor spatiotemporal domains and likely has a large impact on embryonic transcriptional regulatory networks including those that help define the future plant body plan (Figures 5 and 6C) (Nodine and Bartel, 2010; Seefried et al., 2014). To determine how miRNA-mediated cleavage of transcripts encoding transcription factors contributes to embryo morphogenesis, we selected six miRNA:target interactions to study further including miR156/157:SQUAMOSA PROMOTER BINDING PROTEIN-LIKE 10 (SPL10), miR156/157:SPL11, miR160:AUXIN RESPONSE FACTOR 17 (ARF17), miR165/166:PHB, miR167:ARF8 and miR319:TCP4 (Figure 6B and Figure 7A).

**Figure 7.**
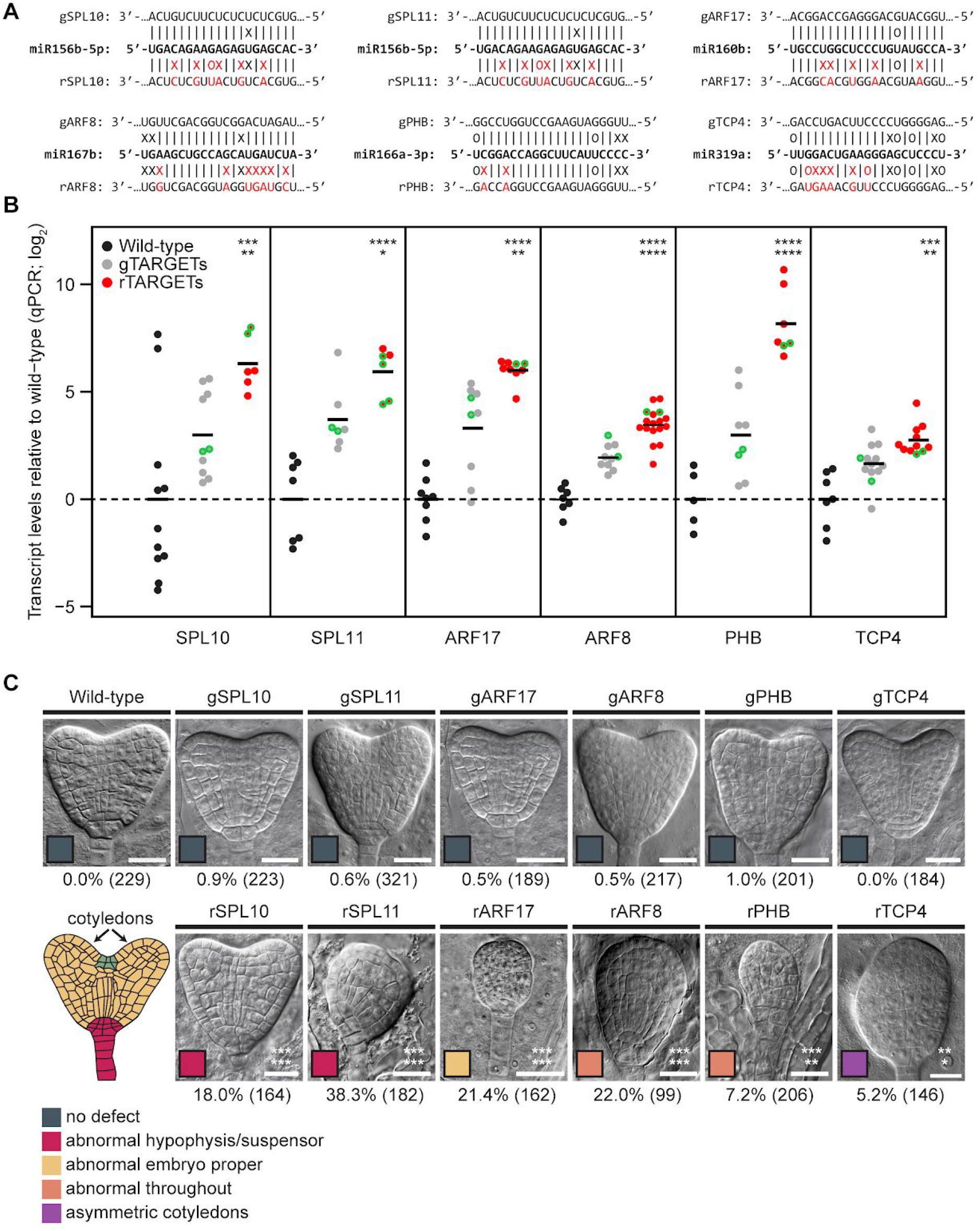
miRNA-Mediated Repression of Transcription Factors is Required for Embryo Morphogenesis. (A) Schematics of six miRNA target sites in transcripts encoding transcription factors selected for mutagenesis. The dominant miRNA isoforms in globular stage embryos for each family are shown, as well as base-pairing interactions with either wild-type target sites (genomic; gTARGET) or miRNA-resistant target sites (resistant; rTARGET) are indicated above and below, respectively. Mutations introduced by site-directed mutagenesis are labelled in red. Watson-Crick base-pairing (I), non-base-pairing (X) and G:U wobbles (O) for each pair are indicated. (B) miRNA target transcript levels based on qRT-PCR on flowers from wild-type plants, or plants expressing either wild-type (gTARGET) or miRNA-resistant (rTARGET) versions of the target transcripts. Values were internally normalized with transcript levels from the eIF4A1 housekeeping gene, and then divided by the levels observed in wild-type plants and log_2_-transformed. Each dot represents the mean of two technical replicates from an independent transgenic line (first generation; T_1_ lines), and are color-coded according to the key. Horizontal bars represent the means between all lines. Asterisks indicate whether the transcript levels observed in rTARGET lines were significantly different compared to either wild-type (top) or gTARGET controls *(bottom)* (two-tailed Student’s *t*-tests; ****, ***, ** and * represent *P*-values < 0.0001, *P*-values < 0.001, *P*-values < 0.01 and *P*-values <0.05, respectively). Points corresponding to lines selected for phenotypic analyses are outlined in green. (C) Representative Nomarski images of embryos derived from crosses between wild-type mothers and fathers that were either wild-type, or expressed transgenic copies of target transcripts containing wild-type (genomic; gTARGET) or abolished (resistant; rTARGET) miRNA binding sites 120 hours after pollination. Genotypes of paternal parents used in the crosses are labelled above each image. The percentages of abnormal embryos for each genotype are shown below each image along with the total number of embryos phenotyped. Asterisks indicate whether the number of abnormal embryos was greater than either wild-type (top) or gTARGET control (bottom) embryos (one-tailed Fisher’s exact test; ***, ** and * represent *P*-values < 0.0001, *P*-values < 0.001 and *P*-values < 0.01, respectively). Scale bars represent 20 μm. A diagram illustrating heart stage embryos is shown *(bottom left)* where different regions are color-coded according to the key, and also correspond to the color-codes in each image which indicate the dominant abnormal phenotype observed. See also Figure S5.

We cloned the target loci, including upstream and downstream intergenic sequences, and performed site-directed mutagenesis to introduce synonymous mutations in the corresponding miRNA target sites in order to abolish miRNA binding and generate transgenic targets resistant to miRNA-mediated cleavage (i.e. resistant targets) (Figure 7A). To control for potential effects unrelated to the disruption of the miRNA binding sites, we also generated constructs whereby the miRNA binding sites were left intact (i.e. genomic targets) (Figure 7A). We transformed the resistant and genomic target constructs into wild-type plants, and selected 6-17 independent transgenic lines for each. Post-embryonic phenotypes of resistant target lines were consistent with previous reports (Mallory et al., 2005; Wu et al., 2009, 2006) (Figure S5A). To characterize these transgenic lines further, we performed quantitative RT-PCR (qRT-PCR) on target transcripts in floral buds from wild-type, or their respective genomic and resistant target lines (Figure 7B). Significantly higher transcript levels were observed in all sets of resistant lines relative to wild-type and corresponding genomic target control lines (*P*-values < 0.001 and 0.05, respectively; *t*-tests) (Figure 7B).

Based on the qRT-PCR analyses, we selected two representative lines for each construct to examine for embryo abnormalities (Figure 7B). Because most of the resistant target lines were sterile and had abnormal flower morphologies (Figure S5B), we crossed wild-type mothers with fathers that were either wild-type, or transgenic miRNA-resistant or control target lines to determine whether specific miRNA/target interactions were required for embryo morphogenesis. More precisely, we fixed and cleared ovules 120 hours after pollination, and inspected the resulting embryo morphologies with Nomarski microscopy. Strikingly, all crosses between wild-type mothers and resistant target lines yielded significantly more abnormal embryos than wild-type mothers crossed with either wild-type or genomic target control line fathers (*P*-values < 0.001 and < 0.01, respectively; Fisher’s exact test) (Figure 7C). None of the genomic control line crosses had significantly more abnormal embryos relative to wild-type (Figure 7C). Moreover, different resistant targets generated distinct embryonic phenotypes including abnormal cell divisions in either basal regions of the embryo (i.e. hypophysis and suspensors) (rSPL10 and rSPL11), embryo propers (rARF17), embryo propers and suspensors (rARF8 and rPHB), or cotyledon growth defects (rTCP4) (Figure 7C).

Importantly, these characteristic phenotypes were reproducible between independent transgenic lines (Figure S5C). Therefore, specific interactions between miRNAs and transcripts encoding transcription factors are morphologically required in a variety of embryonic cell-types.

## DISCUSSION

We developed a sRNA-seq library preparation protocol that can be used to highly enrich for sRNAs, as well as reproducibly and accurately measure their levels from low amounts of total RNA (i.e. ≥1 ng). We expect this method to be useful for profiling sRNA populations from difficult-to-obtain samples including plant and animal reproductive tissues. Here, we used this low-input sRNA-seq method to profile sRNA populations across *Arabidopsis* embryogenesis. In general, the sRNA-seq and nanoPARE datasets reported in this study, as well as the transcriptome datasets produced from the same stages (Hofmann et al., 2019), provide a solid foundation for the characterization of non-coding and coding RNA populations in plant embryos. For example, miRNAs comprise only a fraction of the embryonic sRNA population, and these integrated sRNA-seq, nanoPARE and mRNA-seq datasets will also enable the systematic characterization of additional sRNAs present in early embryos including small interfering RNAs that may be involved in the establishment of epigenetic marks and associated transcriptional gene silencing events.

In this study, we generated a catalog of 354 miRNAs present during embryogenesis and applied our recently developed nanoPARE method to identify 59 high-confidence embryonic miRNA targets. We found high-confidence miRNA-directed cleavage products for only 22/115 miRNAs detected at ≥1 RPM in at least one stage suggesting that many miRNAs may not be directing target cleavage in the stages and conditions examined. Although this could be partially explained by limited sensitivity of the nanoPARE method, our observation that targets detected by nanoPARE, but not those confidently predicted computationally, had globally increased transcript levels in *dcl1-5* embryos suggested that we have identified the majority of cleavage events. Moreover, we detected miRNA-directed cleavage products of all targets with published evidence supporting their existence during *Arabidopsis* embryogenesis as described below. We propose that many of the detected miRNAs function as fail-safes to prevent the aberrant accumulation of target transcripts or have already executed their functions during earlier stages of development. For instance, we were unable to detect targets for any of the five miRNA families that were abundant and enriched at the earliest stages of embryogenesis. These miRNAs decreased dramatically during early embryogenesis and we propose that they function directly after fertilization and prior to the earliest stage sampled (i.e. preglobular or 8-cell/16-cell stage). miRNAs for which targets were not detected may also recognize their targets through mechanisms that are not accounted for by miRNA target computational predictions. For example, miR156/157:AT5G64640, miR158:SPY, and miR159:ATPD sites have relatively poor miRNA:target RNA duplex complementarities and are consequently predicted to be unlikely targets (i.e. Allen scores ≥5). Although future characterization of these embryo-enriched miRNA-directed cleavage events is required to determine their functional relevance, all three cleavage products were robustly detected specifically during embryogenesis suggesting that the full spectrum of miRNA targets remains to be identified even in *Arabidopsis* for well-characterized and highly conserved miRNAs.

As a proof-of-principle of this resource’s utility, we focused on the miRNA-mediated regulation of transcription factors in the current study. We and others demonstrated that miRNAs were required for pattern formation and proper developmental timing of gene expression programs during *Arabidopsis* embryogenesis (Nodine and Bartel, 2010; Seefried et al., 2014; Willmann et al., 2011). Indeed, the more comprehensive *dcl1-5* embryo transcriptome dataset and analyses presented here further supports that miRNAs have a large impact on the embryonic transcriptome including the prevention of precocious expression of genes characteristic of the maturation phase of embryogenesis and related to oil body biogenesis, lipid storage and other seed maturation processes (Hofmann et al., 2019). Because < 5% of the transcripts significantly increased in *dcl1-5* embryos were directly cleaved and repressed by miRNAs, we suggest that the vast majority of mis-regulated genes in *dcl1-5* embryos are downstream of miRNA targets. Interestingly, 30 of the 59 embryonic miRNA high-confidence targets identified encode transcription factors and their de-repression in miRNA-deficient *dcl1-5* embryos, along with associated mis-regulated downstream transcriptional cascades, may largely explain why thousands of transcripts have different levels in *dcl1-5* compared to wild-type embryos.

Together with previous studies, our results indicate that multiple miRNAs are required for embryo morphogenesis and pattern formation. We previously demonstrated that miR156 prevents the accumulation of SPL transcription factors and the resulting expression of maturation phase genes during early embryogenesis (Nodine and Bartel, 2010). Although, decreased SPL10/11 levels could suppress miRNA-deficient *dcl1-5* phenotypes, abolishing miR156:SPL10/11 interactions were not sufficient to phenocopy *dcl1-5* embryos. This suggests that additional miRNA:target interactions are required for embryonic pattern formation. Consistent with this idea, the phenotypes of embryos with *mir160a* loss-of-function mutations (Liu et al., 2010) or expressing miR160-resistant ARF17 transgenes (Figure 7C) indicated that miR160-mediated repression of the ARF17 transcription factor is required for proper cell division patterns in the embryo proper. Seedling (Palatnik et al., 2003) and embryo phenotypes of plants expressing miR319-resistant TCP4 transcription factor transgenes (Figure 7C) demonstrated that miR319:TCP4 interactions are required for proper cotyledon formation. Previous results, as well as our characterization of miR165/166 and their target HD-ZIP transcript localization patterns, indicated that miR165/166 helps define HD-ZIP transcription factor localization domains in early embryos (McConnell et al., 2001; Miyashima et al., 2013; Smith and Long, 2010) (Figures 3, 4 and 6). Consistent with miR165/166-directed spatial restriction of HD-ZIP transcription factors being required for embryonic pattern formation, embryos expressing the miR165/166-resistant PHB transcription factor exhibited abnormal divisions (Figure 7C). Interestingly, the morphological defects observed in rARF8 embryos are consistent with previous reports that miR167:ARF8 interactions are required in the maternal sporophytic tissues for proper embryogenesis (Wu et al., 2006; Yao et al., 2019). However, miR167 and miR167:ARF8 cleavage products were nearly eliminated in *dcl1-5* embryos derived from self-fertilized *dcl1-5/+* plants (Table S2, Figures 3A and 4B), suggesting that the activities of zygotically produced miR167 are required for embryo morphogenesis. Our identification of miR170/171- and miR394-directed cleavage of transcripts respectively encoding either the HAM1 transcription factor or LCR F-box protein is also consistent with previous reports (Knauer et al., 2013; Takanashi et al., 2018). Altogether, these data support a model whereby the post-transcriptional regulation of transcription factor gene-regulatory networks by several miRNAs is critically important for the establishment of the plant body plan during early embryogenesis. The resources and phenotypes described in this study therefore provide multiple entry points to further characterize how the miRNA-mediated repression of transcripts, including those encoding for transcription factors, contributes to the initial cellular differentiation events operating at the beginning of plant life.

## ACKNOWLEDGEMENTS

We thank the VBCF NGS Unit for sequencing sRNA-seq, nanoPARE and mRNA-seq libraries and the VBCF Plant Sciences Facility for plant growth chamber access. This work was supported by funding from the European Research Council under the European Union’s Horizon 2020 research and innovation program (Grant 637888 to M.D.N.) and the DK Graduate Program in RNA Biology (DK-RNA) sponsored by the Austrian Science Fund (FWF, DK W 1207-B09).

## AUTHOR CONTRIBUTIONS

Conceptualization, M.D.N.; Methodology, M.D.N., A.P., M.J.K. and M.M.; Software, M.D.N. and M.A.S.; Formal Analysis, M.D.N. and M.A.S.; Investigation, M.D.N., A.P., M.J.K. and M.M.; Writing – Original Draft, M.D.N.; Writing – Review & Editing, M.D.N., M.A.S., M.J.K. and A.P.; Visualization, M.D.N., M.A.S. and A.P.; Supervision, M.D.N.; Funding Acquisition, M.D.N.

## DECLARATION OF INTERESTS

The authors declare that they have no conflicts of interest.

## METHODS

### Plant Material, Growth Conditions and RNA isolation

The *dcl1-5* (McElver et al., 2001) and *xrn4-5* (Souret et al., 2004) alleles were in Col-0 accession backgrounds, and together with Col-0 were grown in a climate-controlled growth chamber with 20°C-22°C temperature and 16h light/8h dark cycle. Embryos were dissected and total RNA was extracted at a similar time of day (13:00-17:00) as described previously (Hofmann et al., 2019). Except for the bent-cotyledon stage samples, all other total RNA samples pooled from 50 Col-0 embryos were used to generate mRNA-seq datasets (Hofmann et al., 2019), and the sRNA-seq and nanoPARE datasets reported in this study. Total RNA from *xrn4-5* roots and shoots were isolated as previously described (Schon et al., 2018).

### Low-input sRNA-seq

RNAs of 18-30-nt lengths were purified from ≥80% of each sample’s total RNA (from 50 pooled embryos) using denaturing polyacrylamide-urea gels as described previously (Grimson et al., 2008). Size-selected sRNAs were precipitated overnight at −20°C with 2.5× volumes of ice-cold 100% ethanol and 1 μl of GlycoBlue (Thermo Fisher), and resuspended in 7.5 μl of nuclease-free water. This was used as input for the NEBNext Multiplex Small RNA Library Prep Set for Illumina kit (NEB #E7300) according to the manufacturer’s recommendations with the following modifications. Adapters used for 3’ and 5’ ligations to sRNAs, and SR RT primers to generate sRNA cDNAs, were diluted to 25% of the amounts recommended for ≥500 ng of total RNA. Various numbers of PCR cycles were used to amplify cDNAs: 14, 16, 18 and 20 PCR cycles for early heart and later staged samples, and 18, 20, 22 and 24 PCR cycles for globular and earlier staged samples. Final amplicons of 137-149-bp (corresponding to 18–30-nt sRNAs) were size-selected and purified on a 90% formamide/8% acrylamide gel. Fluorescence intensities of amplicons were examined across the PCR cycles, 137–149-bp products with non-saturated signals were gel-purified, and after DNA precipitation pellets were resuspended with 15 μl of Elution Buffer (Qiagen). To control for library quality, sRNA-seq libraries were examined on an Agilent DNA HS Bioanalyzer Chip and those with the expected size range were sequenced on an Illumina HiSeq 2500 instrument in 50 base single-end mode (Table S1).

Cutadapt (Martin, 2011) was used to trim adapter sequences from sRNA-seq reads and 18–30-base sequences that contained an adapter were retained. The trimmed sequences were aligned to the *Arabidopsis thaliana* TAIR10 genome (Lamesch et al., 2012) with STAR (Dobin et al., 2013) requiring no mismatches and allowing ≤100 multiple end-to-end alignments. Resulting SAM files were then processed with the readmapIO.py script to re-assign multimappers with a “rich-get-richer” algorithm as previously described (Schon et al., 2018). Output bedFiles were sorted, condensed and normalized for total genome-matching reads. The BEDtools *map* function (Quinlan and Hall, 2010) was then used to quantify the number of reads mapping to the same strand and overlapping ≥80% of mature miRNAs as annotated in TAIR10 and miRBase21 (Kozomara et al., 2019). Statistical analyses and associated figures were generated with the R statistical computing package (R Core Team, 2018).

### nanoPARE and mRNA-seq

Col-0 mRNA-seq data were downloaded from the NCBI Gene Expression Omnibus (GEO; https://www.ncbi.nlm.nih.gov/geo/; accession number GSE121236). An identical procedure was used to generate Smart-seq2 libraries (Picelli et al., 2013) from *dcl1-5* embryos selected from self-fertilized *dcl1-5/+* plants based on their abnormal morphologies, and were sequenced on an Illumina HiSeq 2500 instrument in 50 base paired-end mode (Table S1). Transcriptome analyses were as described (Hofmann et al., 2019) except that TAIR10 transposable element gene models were also included in the Kallisto-based pseudo-alignments (Bray et al., 2016). DESeq2 (Love et al., 2014) with default settings was used to compute *P*-values for the wild-type and *dcl1-5* transcriptome comparisons.

nanoPARE data from floral buds and floral organs were downloaded from the NCBI GEO (accession number GSE112869). nanoPARE libraries presented in this study were generated as previously described (Schon et al., 2018) with the following exceptions. For all embryonic samples other than those from the bent-cotyledon stage, the same cDNA pools used in our previous transcriptome analysis (Hofmann et al., 2019) were also used as input for nanoPARE library preparation. nanoPARE libraries were sequenced on an Illumina Hi-Seq 2500 instrument in 50 base single-end mode (Table S1). Analysis of nanoPARE data was as described (Schon et al., 2018) except that all capped features identified in the embryonic series were merged with those from published floral bud samples (Schon et al., 2018), and used to mask capped features from the transcript-level bedGraph files. Additionally, we used *EndCut* (Schon et al., 2018) to test for significant target sites for 164 miRNAs detected ≥1 RPM in at least one embryonic stage. nanoPARE libraries from all post-embryonic tissues were analyzed in an identical manner.

### RNA in situ Hybridizations

miRNA in situs on embryo sections were performed based on a whole-mount in situ hybridization method (Ghosh Dastidar et al., 2016). Sample preparation leading up to probe hybridization was as described (Nodine et al., 2007) except that a LOGOS Microwave Hybrid Tissue Processor (Milestone Medical) was used for tissue embedding and samples were fixed with EDC solution (0.16 M N-(3-Dimethylaminopropyl)-N’-ethylcarbodiimide hydrochloride in Methylimidazole-NaCl) after the proteinase K digestion step as follows. First, slides with adhered embryo sections were transferred to 1× PBS and washed 2×, and then incubated in a staining dish containing freshly prepared methylimidazole-NaCl for 10 minutes at room temperature (2×). The slides were then transferred to EDC solution and incubated for 2 hours at 60°C, and subsequently washed 2× in 1× PBS for 5 minutes each prior to probe hybridization. Dual DIG-labelled LNA-modified oligos antisense to miR124, miR156a-f, miR159a, miR160a-c or miR166a-f isoforms were used at 20 nM final concentrations (Table S5), and the rest of the probe hybridization procedure, as well as subsequent washing, antibody and colorimetric reactions were as described (Nodine et al., 2007). Slides were imaged on an automated Pannoramic SCAN 150 slide scanner (3DHISTECH) and collected with the associated Pannoramic Viewer software. Images of ≥50 embryos from >5 independent sets of experiments were recorded.

mRNA in situs were as previously described (Nodine et al., 2007). Probes antisense to CNA, PHB and PHV were generated from cDNAs by introducing T7 promoters with PCR as described previously (Hejátko et al., 2006) (Table S5).

### Generation of Transgenic Lines

Control genomic and miRNA-resistant SPL10 and SPL11 constructs were generated as previously described (Nodine and Bartel, 2010). For control genomic ARF17 (gARF17), PHB (gPHB) and TCP4 (gTCP4) transgenic constructs, target loci including upstream and downstream intergenic sequences were PCR amplified from Col-0 genomic DNA with primers containing overhangs for subsequent Gibson assembly. miR160-resistant ARF17 (rARF17), miR165/166-resistant PHB (rPHB) and miR319-resistant TCP4 (rTCP4) constructs were amplified as two separate fragments with overlaps to introduce specific mutations in the corresponding miRNA target sites. The backbones of the MultiSite-Gateway destination vectors pAlligatorG43 and pAlligatorR43 (Kawashima et al., 2014) were amplified for subsequent Gibson assembly, and genomic and resistant ARF17, PHB and TCP4 plant transformation constructs were generated by Gibson Assembly (NEB) using the pAlligatorG43/R43 backbone and the target PCR fragments described above. For the control genomic ARF8 transgenic construct (gARF8), the ARF8 locus including upstream and downstream intergenic sequences was PCR amplified from Col-0 genomic DNA and cloned into the pENTR/D-TOPO Gateway vector (Thermo Fisher). The miR167-resistant ARF8 construct (rARF8) was generated by PCR site-directed mutagenesis (NEB) of the gARF8 entry clone. Final plant transformation constructs were generated by Gateway LR reactions (Thermo Fisher) with pENTR-gARF8 or pENTR-rARF8, pDONR-L4R1-empty and pDONR-R2L3-empty, and the Gateway destination vector pAlligatorR43 (RFP). Constructs were transformed into Col-0 via the Agrobacterium floral dip method (Clough and Bent, 1998). Transformants were selected based on GFP or RFP selection marker fluorescence from pAlligatorG43/R43 (Bensmihen et al., 2004; Kawashima et al., 2014).

### qRT-PCR Analysis

Two clusters of floral buds were pooled from 8-week old plants, snap-frozen in liquid nitrogen, homogenized using a Mixer Mill MM 400 (Retsch) for 30 seconds with max amplitude and resuspended in 200 μl TRIzol (Life Technologies). Total RNA was extracted using the Direct-zol kit (Zymo Research) according to the manufacturer’s instructions, and DNaseI treatment was performed on-column. Total RNA qualities and quantities were determined with an Agilent Fragment Analyzer (AATI) using the standard RNA sensitivity kit (DNF-471). 200 ng of total RNA samples with RQN values >6.0 were used for cDNA synthesis together with the Oligo d(T)_18_ mRNA Primer (NEB) and SuperScript III Reverse Transcriptase (Thermo Fisher). cDNA was diluted 10× with nuclease-free water and 2 μl was used as a template for the qRT-PCR. qRT-PCR was performed on a LightCycler 96 Instrument (Roche) using gene-specific and control elF4A primes (Table S5), and Fast SYBR™ Green Master Mix (Roche). Ct values were obtained by the LightCycler 96 software and relative quantification of transcripts (ΔΔCt values) were calculated with an in-house R script. For each genotype, 6-17 individual first-generation transgenic (T_1_) lines were analyzed in technical duplicates.

### Genetic Crosses and Microscopy

Stage 12 Col-0 floral buds were emasculated and pistils were left overnight before hand-pollinating with at least two independently generated second-generation transgenic (T_2_) lines that had representative transcript levels based on the qRT-PCR experiments described above. Siliques were harvested 120 hours after pollination, and ovules were fixed and cleared in a solution composed of 8 g chloral hydrate, 1 ml water and 1 ml glycerol as described previously (Ohad et al., 1996). Embryos were examined with Nomarski optics on a ZEISS Axio Observer Z1 with a sCMOS camera. Images were acquired using ZEISS ZEN (blue edition) imaging software and analyzed using ImageJ/Fiji processing software. To minimize potential bias, all microscopy images were examined by a person that did not record the images, and phenotypes were recorded prior to revealing sample identities.

## SUPPLEMENTAL INFORMATION

**Table S1 Summary of High-Throughput Datasets Generated In or Reanalyzed for This Study, Related to Figures 2, 4 and 5**

**Table S2 Levels of miRNAs Detected During Embryogenesis, Related to Figure 3**

**Table S3 Predicted miRNA Cleavage Sites Detected in nanoPARE Datasets, Related to Figure 4**

**Table S4 Normalized Transcript Levels in Wild-type and miRNA-Deficient *dcl1-5* Globular Embryos, Related to Figure 5**

