## Supplemental File for "MicroRNA Dynamics and Functions During *Arabidopsis* Embryogenesis"

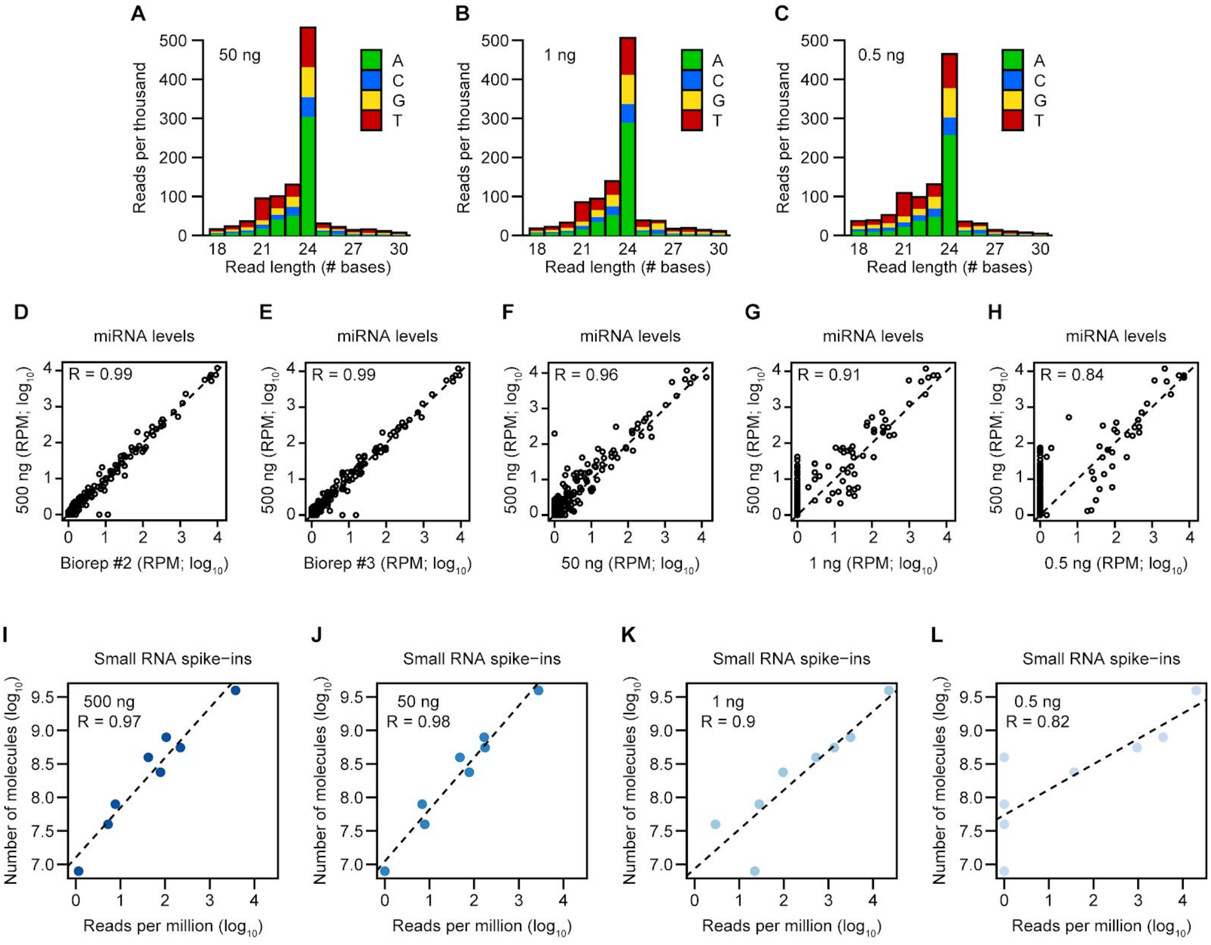


**Figure S1.** Establishment of Low-input Small RNA Sequencing Method, Related to Figure 1

(A to C) Stacked bar charts of normalized sRNA levels (reads per thousand genome-mapping reads) across different nucleotide (nt) lengths in libraries generated with either 50 ng (A), 1 ng (B) or 0.5 ng **(C)** of total RNA isolated from bent cotyledon stage embryos. Colors indicate proportions of sRNA-seq reads that begin with various bases as indicated in key.

(D to H) Scatter plots of miRNA family levels in sRNA-seq libraries generated from either 500 ng (biological replicate #2) (D), 500 ng (biological replicate #3) (E), 50 ng (F), 1 ng (G) or 0.5 ng (H) compared to 500 ng of total RNA (biological replicate #1). sRNA levels were normalized for reads per million genome-mapping reads (RPM) and $log_{10}$-transformed. Pearson's *R* values are indicated, as well as a dashed line with an intercept of 0 and slope of 1.

(I to L) Scatter plots of relative sRNA spike-in levels (RPM; $log_{10}$) compared to the absolute number of sRNA spike-in molecules ($log_{10}$) added during RNA isolation for a sRNA-seq library generated from either 500 ng (I), 50 ng (J), 1 ng (K) or 0.5 ng (L) of total RNA. Pearson's *R* values are shown, and the dashed lines represent linear models derived from the plotted data points.

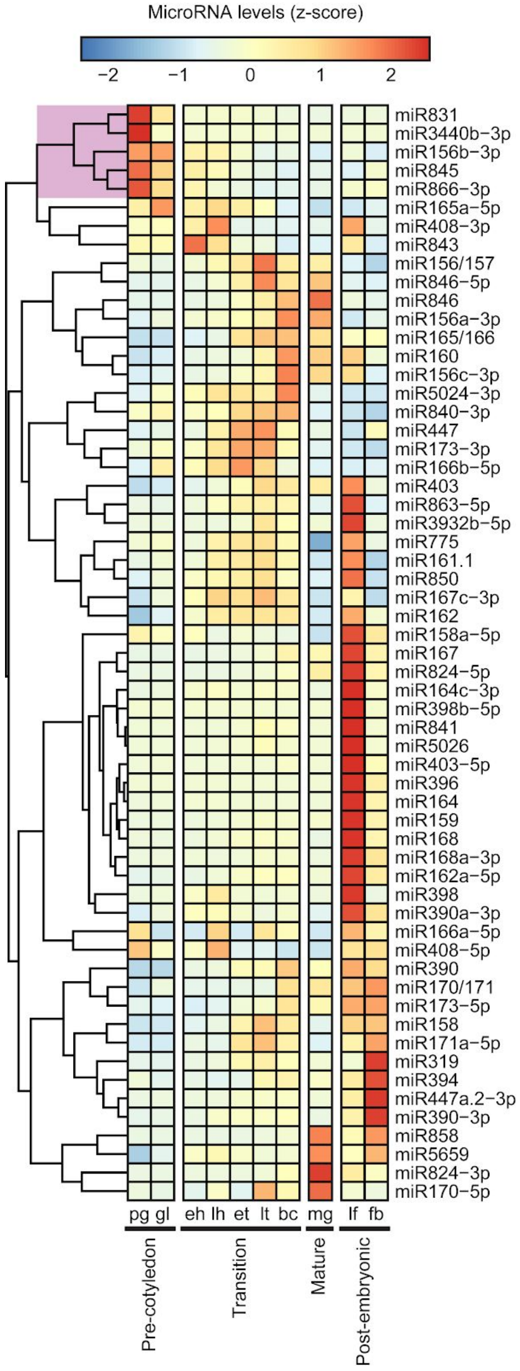


**Figure S2.** Embryo-Enriched miRNAs, Related to Figure 3

Heat map illustrating the relative levels of miRNA families across embryogenesis, leaves and floral buds. miRNA families with ≥10 mean RPM in at least one embryonic stage are shown, and colors represent z-scores for each individual miRNA family according to the key. Three major phases of embryo development are indicated at the bottom and individual columns are labelled according to stage: pg, preglobular; gl, globular; eh, early heart; lh, late heart; et, early torpedo; lt, late torpedo; bc, bent cotyledon; mg, mature green; lf, leaves; fb, unopened floral buds. The dendrogram clade color-coded in violet indicates the five miRNA families enriched in early embryos.

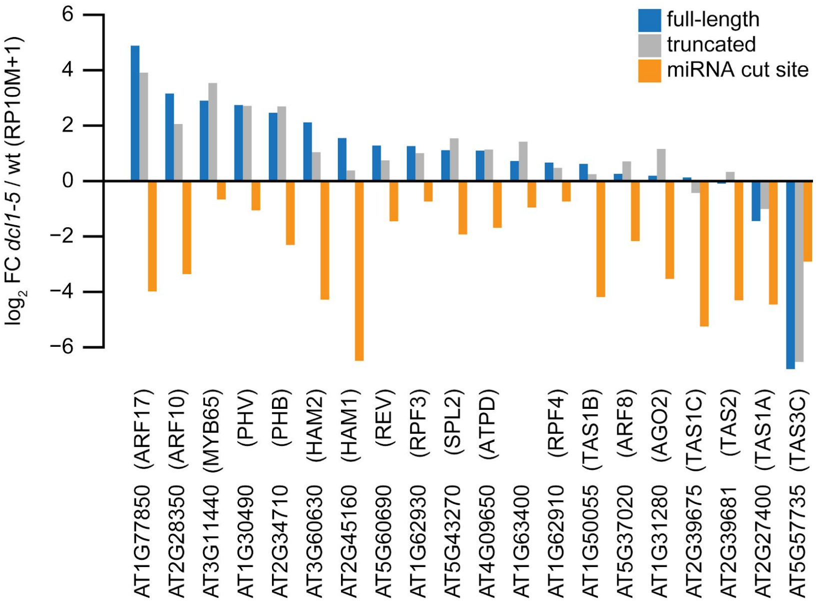


**Figure S3.** mRNA 5′ Ends of miRNA Targets in *dcl1-5* Mutant Embryos, Related to Figure 4

Comparison of the abundance of nanoPARE read 5′ ends mapping to different positions within all 20 high-confidence miRNA target transcripts identified in wild-type globular embryos. Each gene was subdivided into three regions: blue, annotated transcription start sites identified with nanoPARE (Schon et al. 2018); orange, positions 9 and 10 of the miRNA:target site; gray, all other exonic positions in the gene. Y-axis represents $\log_2$ fold change of the mean abundance of each gene feature in globular-stage *dcl1-5* mutant embryos compared to wild-type embryos of the same stage (reads per ten million genome-matching reads, RP10M).

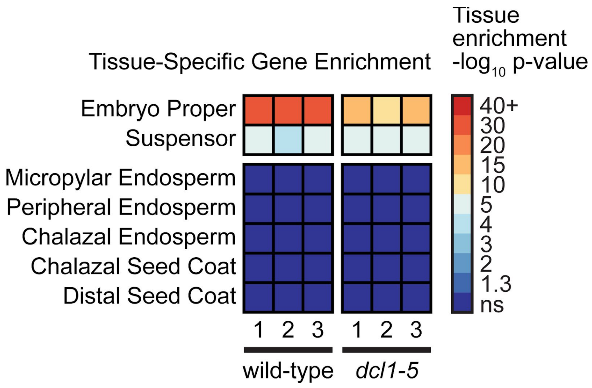


**Figure S4.** Tissue-Enrichment Test of Wild-Type and *dcl1-5* Mutant Embryo Transcriptomes, Related to Figure 5
Statistical enrichment of seven distinct seed tissue types inferred through the expression of tissue-enriched gene sets using the tissue-enrichment test (Schon and Nodine 2017) with default parameters. The three wild-type replicates are globular-stage mRNA-seq samples from GEO series GSE121236; *dcl1-5* replicates were generated for this study.

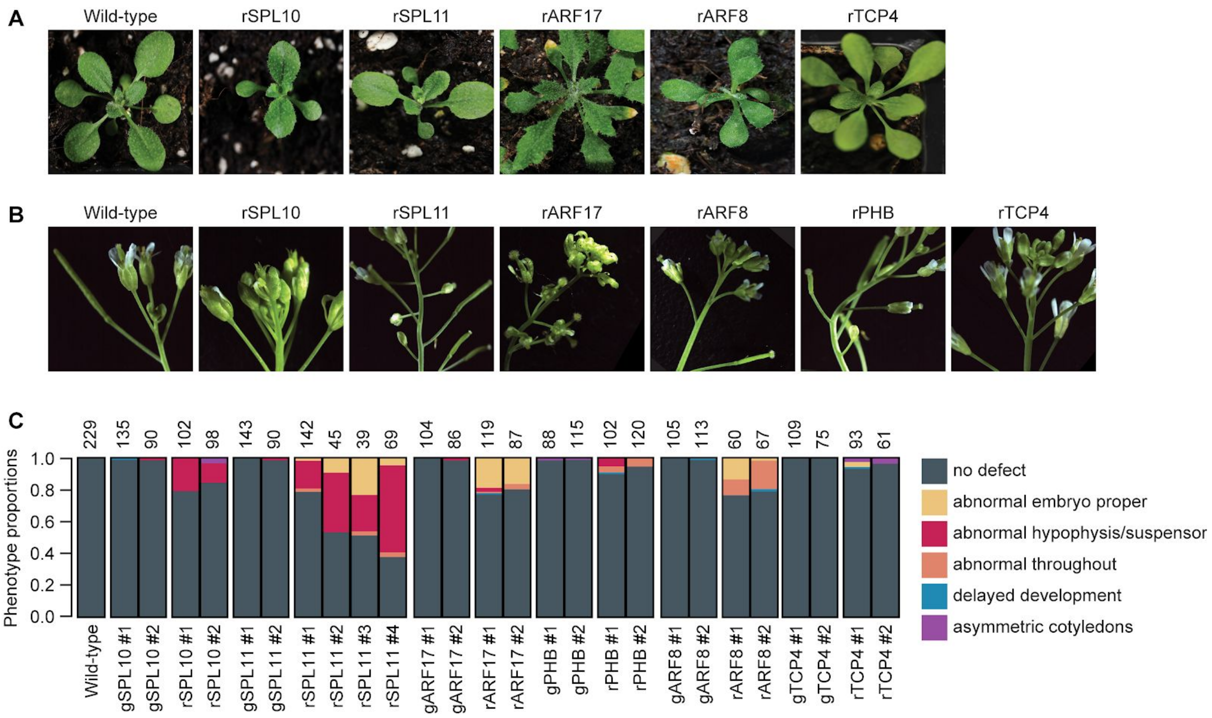


**Figure S5.** Post-Embryonic Phenotypes of Plants Expressing miRNA-Resistant Targets and Quantification Details, Related to Figure 7

(A-B) Representative images of vegetative (A) or flowering (B) plants (A) expressing miRNA-resistant targets. Genotypes are indicated above each panel.

(C) Stacked bar plot illustrating the proportions of phenotypes observed for embryos derived from crosses between wild-type mothers and fathers that were either wild-type, or expressed transgenic copies of target transcripts containing wild-type (genomic; gTARGET) or abolished (resistant; rTARGET) miRNA binding sites 120 hours after pollination. Paternal genotypes used in the crosses, including transgenic line numbers, are labelled below. Numbers above each bar denote how many embryos were examined. Phenotypes are color-coded according to the legend.

**Table S5.** Oligonucleotides Used in This Study

| Name | Sequence | General purpose | Specific purpose |
|---|---|---|---|
| miR124-AS-LNA | /5DIGN/ACTGATA+TC+AG+CTC+AGTAG GCAC/3DIG_N/ | miRNA in situs | Negative control; antisense to animal-specific miR124 (+ indicates LNA position, /5DIGN/ and /5DIGN/ represents digoxigenin labels on the 5′ and 3′ ends) |
| miR156-AS-LNA | /5DIGN/GTGCT+CACT+CT+CTTCTG+TCA /3DIG_N/ | miRNA in situs | Antisense to miR156a-f isoforms (+ indicates LNA position, /5DIGN/ and /5DIGN/ represents digoxigenin labels on the 5′ and 3′ ends) |
| miR159-AS-LNA | /5DIGN/TAGAG+CT+CC+CTT+CAATC+CA AA/3DIG_N/ | miRNA in situs | Antisense to miR159a isoform (+ indicates LNA position, /5DIGN/ and /5DIGN/ represents digoxigenin labels on the 5′ and 3′ ends) |
| miR160-AS-LNA | /5DIGN/GGCATA+CAGG+GAG+CCAGG+ CAC/3DIG_N/ | miRNA in situs | Antisense to miR160a-c isoforms (+ indicates LNA position, /5DIGN/ and /5DIGN/ represents digoxigenin labels on the 5′ and 3′ ends) |
| miR166-AS-LNA | /5DIGN/GGGGAA+TGAA+GC+CTGGTC+C GA/3DIG_N/ | miRNA in situs | Antisense to miR166a-f isoforms (+ indicates LNA position, /5DIGN/ and /5DIGN/ represents digoxigenin labels on the 5′ and 3′ ends) |
| PHB F10 | GTAGCGATGGTGCAGAGGATGT | mRNA in situs | Forward primer for amplifying PHB amplicon from cDNA |
| PHB R10 | CGAACGACCAATTCACGAACAT | mRNA in situs | Reverse primer for amplifying PHB amplicon from cDNA |
| PHB F12 | GTAGCGATGGTGCAGAGGATGTTACTG | mRNA in situs | Forward primer for incorporation of T7 site for PHB antisense probe generation |
| PHB R12T7 | CCAAGCTTCTAATACGACTCACTATAGG GAGACGAACGACCAATTCACGAAC | mRNA in situs | Reverse primer for incorporation of T7 site for PHB antisense probe generation |
| PHB F11T7 | CCAAGCTTCTAATACGACTCACTATAGG GAGAGTAGCGATGGTGCAGAGGAT | mRNA in situs | Forward primer for incorporation of T7 site for PHB sense probe generation |
| PHB R11 | CGAACGACCAATTCACGAAC | mRNA in situs | Reverse primer for incorporation of T7 site for PHB sense probe generation |
| CNA_F1 | TTCAAAGGCAACTGGAACCG | mRNA in situs | Forward primer for amplifying CNA amplicon from cDNA and for CNA antisense probe generation |
| CNA_R1 | CCGCACAGGTTCCTACAGCA | mRNA in situs | Reverse primer for amplifying CNA amplicon from cDNA |
| CNA_AS_R1T7 | CCAAGCTTCTAATACGACTCACTATAGG GAGAAGCAAGTGGAAGTATAACCT | mRNA in situs | Reverse primer for incorporation of T7 site for CNA antisense probe generation |
| PHV_F1 | GGTCGCTGAAATCCTCAAAG | mRNA in situs | Forward primer for amplifying PHV amplicon from cDNA |
| PHV_R1 | TTCGGATTTGTTTTTTGGTC | mRNA in situs | Reverse primer for amplifying PHV amplicon from cDNA |
| PHV_AS_F1 | TCGTCCATCTTGGTTCCGTG | mRNA in situs | Forward primer for incorporation of T7 site for PHV antisense probe generation |
| PHV_AS_R1T7 | CCAAGCTTCTAATACGACTCACTATAGG GAGAATTTTCATCAACGCCGCTAC | mRNA in situs | Reverse primer for incorporation of T7 site for PHV antisense probe generation |
| pAlligatorR/G43-F1 | CTGCAGATCGTTCAAACATTTG | Cloning | Forward primer for generating the backbone of pAlligatorG43/R43 (use for g/mARF17,CAN, PHB, TCP4) |
| pAlligatorR/G43-R1 | CTGCAGGTCGACCATAGTG | Cloning | Reverse primer for generating the backbone of pAlligatorG43/R43 (use for g/mARF17,CAN, PHB, TCP4) |
| pAlligatorR/G43-R2 | ATAGCTTGGCGTAATCATGG | Cloning | Alternative reverse primer for generating the backbone of pAlligatorG43/R43 (use for g/mPHB) |
| g/mARF17-F1 | ACACAACATATCCAGTCACTATGGTCGA CCTGCAGTGTCTTTTGTTTTTAGGTTTTT TTTTTAAC | Cloning | Forward primer for genomic ARF17 amplification and gibson cloning into pAlligatorG43 /pAlligatorR43 destination vector |
| g/mARF17-R1 | GAAACTTTATTGCCAAATGTTTGAACGAT CTGCAGTTTATTTAGTATTATTTGCTCTG TTTG | Cloning | Reverse primer for genomic ARF17 amplification and gibson cloning into pAlligatorG43 /pAlligatorR43 destination vector |
| rARF17-F2 | CTGGAATGCAAGGTGCACGGCAATATGA TTTTGGGTC | Cloning | Forward primer for amplification of rARF17 gibson piece 2 (with g/mARF17-R1) |
| rARF17-R2 | ACCCAAAATCATATTGCCGTGCACCTTG CATTCCAG | Cloning | Reverse primer for amplification of rARF17 gibson piece 1 (with g/mARF17-F1) |
| gARF8-TOPO-F | CACCTCTCCAAGTGATACACTC | Cloning | Forward primer for genomic ARF8 amplification and cloning into pENTR/D-TOPO |
| gARF8-TOPO-R | TAAGTCTGATGTGTGTGCA | Cloning | Reverse primer for genomic ARF8 amplification and cloning into pENTR/D-TOPO |
| ARF8-SDM-F | CCGGTTGTTACGGAAAATACAAAAACAA C | Cloning | Forward site-directed mutagenesis primer to generate rARF8 |
| ARF8-SDM-R | GGCCTGATTCCATTGGAATCATCG | Cloning | Reverse site-directed mutagenesis primer to generate rARF8 |
| g/mPHB-F1 | AAACAGCTATGACCATGATTACGCCAAG CTATTGGAGGGAAGAGGCTACAAAG | Cloning | Forward primer for genomic PHB amplification and gibson cloning into pAlligatorG43 /pAlligatorR43 destination vector |
| g/mPHB-R1 | ACTTTATTGCCAAATGTTTGAACGATCTG CAGTTGTCCGAGCATTGATTTTGTAC | Cloning | Reverse primer for genomic PHB amplification and gibson cloning into pAlligatorG43 /pAlligatorR43 destination vector |
| rPHB-F2 | AATAGAATCTGGTCCAGGCTACACCAGC AATGAAG | Cloning | Forward primer for amplification of rPHB gibson piece 2 (with g/mPHB-R1) |
| rPHB-R2 | CATTGCTGGTGTAGCCTGGACCAGATTC TATTGGC | Cloning | Reverse primer for amplification of rPHB gibson piece 1 (with g/mPHB-F1) |
| g/mTCP4-F1 | ACACAACATATCCAGTCACTATGGTCGA CCTGCAGCATTTTGATGAGGCGTATATA TATACATTTAATTAATATTG | Cloning | Forward primer for genomic TCP4 amplification and gibson cloning into pAlligatorG43 /pAlligatorR43 destination vector |
| g/mTCP4-R1 | GAAACTTTATTGCCAAATGTTTGAACGAT CTGCAGATATGATCTTTGTGTCATGACT | Cloning | Reverse primer for genomic TCP4 amplification and gibson cloning into pAlligatorG43 /pAlligatorR43 destination vector |

| | | | |
|---|---|---|---|
| | C | | |
| rTCP4-F2 | GGTCCCTTGCAAAGTAGCTACAGTCCCATGATCCGTG | Cloning | Forward primer for amplification of rTCP4 gibson piece 2 (with g/mTCP4-R1) |
| rTCP4-R2 | ACGGATCATGGGACTGTAGCTACTTTGCAAGGGACC | Cloning | Reverse primer for amplification of rTCP4 gibson piece 1 (with g/mTCP4-F1) |
| eIF4A1 RTF | TGCAAGGCACTCTTTGATCTGATTT | qRT-PCR | Forward primer for detection of the housekeeping gene eIF4A1 |
| eIF4A1 RTR | GAGATATGTTCGTAGCTGGGAGAGAGAG | qRT-PCR | Reverse primer for detection of the housekeeping gene eIF4A1 |
| SPL10 RTF | TCAGGAGGCCTCCATGAATCTCA | qRT-PCR | Forward primer for SPL10 detection |
| SPL10 RTR | GGCCACGGGAGTGTGTTTGAT | qRT-PCR | Reverse primer for SPL10 detection |
| SPL11 RTF | CCAACCACATGTGCAGCCATTT | qRT-PCR | Forward primer for SPL11 detection |
| SPL11 RTR | GAACAGAGTAGAGAAAATGGCTGCA | qRT-PCR | Reverse primer for SPL11 detection |
| PHB RTF | GCTAGACAAGACCCTTGACGAACCT | qRT-PCR | Forward primer for PHB detection |
| PHB RTR | TCCCATGCTTGACGCACATACTC | qRT-PCR | Reverse primer for PHB detection |
| ARF8 RTF | CATGCAGATGTTGAGACGGATGAAG | qRT-PCR | Forward primer for ARF8 detection |
| ARF8 RTR | TTACTCGGTATCCCCAACTCAATCG | qRT-PCR | Reverse primer for ARF8 detection |
| ARF17 RTF | GTGCAGCAGCACCTGATCCAAG | qRT-PCR | Forward primer for ARF17 detection |
| ARF17 RTR | GGAGGATTTCCTCCAATGAATCCGG | qRT-PCR | Reverse primer for ARF17 detection |
| TCP4 RTF | CCAGTTCTTGGCCAAAGCCAAC | qRT-PCR | Forward primer for TCP4 detection |
| TCP4 RTR | ATGGTGGTGGTTGAGATCGTCG | qRT-PCR | Reverse primer for TCP4 detection |
